# Estimating the Explainable Variance of EEG Responses to Natural Speech

**DOI:** 10.64898/2026.07.02.736170

**Authors:** Jin Dou, Edmund C. Lalor

## Abstract

Substantial progress has been made in recent years on understanding how the human brain parses and processes natural speech. Much of this progress has been based on modeling how brain activity relates to the different acoustic and linguistic features of speech. By fitting and testing models based on those features, one can test hypotheses about the kinds of computations and representations the brain uses to convert speech sounds into understanding. While much of this work has focused on modeling BOLD activity using functional neuroimaging or intracranially recorded electrophysiological signals, the approach has also proven useful with MEG and EEG. Indeed, noninvasive EEG has certain advantages for studying speech processing in terms of translational research and application. Research over the last decade or so has shown that EEG can be successfully modeled based on numerous acoustic, linguistic, and paralinguistic speech features. However, an important unanswered question hangs over all of this work: namely, what constitutes a *good* model of EEG responses to natural speech? Or, to put it another way, how much variance in EEG recorded during natural speech listening is explainable as having derived from that speech input? The present study aims to tackle this issue. We do so under the assumption that the best model for a person’s EEG response to natural speech is a set of EEG responses from other people listening to the same speech. Using this assumption, we construct inter-subject models using EEG from 19 healthy adult native speakers of English who all listened to the same audiobook. The model for each subject involves predicting their EEG data using (dimensionality-reduced) EEG from different numbers of other subjects and then extrapolating to estimate the total explainable variance in the target individual’s response to speech. Following this, we show that linear models (temporal response functions) based on several commonly used acoustic and linguistic speech features can predict most – but importantly not all – of the estimated total explainable variance in EEG responses across subjects.

## Introduction

How the human brain so effortlessly processes natural speech remains incompletely understood. In recent years, one useful approach to studying this issue involves estimating linear encoding models – sometimes known as temporal response functions (TRFs) – that allow one to model brain responses to speech in terms of various acoustic and linguistic features of that speech (Anderson et al., 2024; Brodbeck et al., 2018; Broderick et al., 2018; Synigal et al., 2023). Typically, such models are fit based on the chosen speech features (e.g., the speech envelope, the lexical surprisal of words) using a subset of one’s data and then tested by assessing how well they can account for (or “predict”) brain responses to speech in left out trials. However, interpreting the predictive accuracy of such models is complicated by at least two issues. First, large inter-individual differences in the signal-to-noise ratio (SNR) of brain responses to speech make it difficult to compare the success of models across subjects based on absolute prediction accuracy. For example, in the case of scalp EEG, differences in cortical folding and/or skull thickness likely contribute to what can be large inter-individual differences in how strongly EEG signals track the dynamics of natural speech (Ahlfors et al., 2010; Antonakakis et al., 2020; Gaudin et al., 2024; Pizzagalli et al., 2020). As such, a “low” absolute correlation (e.g., Pearson’s *r*) between an individual’s real and predicted EEG does not necessarily indicate that the model is bad. Rather, it may simply be that that person’s scalp EEG responses to speech are weak relative to the rest of their ongoing EEG. Second, when comparing how well we can model brain responses to speech based on different combinations of speech and language features, we do not know what constitutes optimal model performance. Adding additional speech and linguistic features may improve a model of brain activity – but how much more brain activity needs to be explained? And, thus, what other speech or linguistic features (or what computations based on those features) might need to be considered in future models? Both of these issues – inter-subject variability and knowledge of optimal model performance – would be addressed if it were possible to estimate the maximum explainable variance in each individual’s brain responses to natural speech.

A common approach to estimating the explainable variance in brain responses to stimuli involves computing correlations between responses to repeated presentations of the same stimulus (Keshishian et al., 2020; Schoppe et al., 2016). This provides a measurement of how consistent the neural activity is across repetitions (Norman-Haignere et al., 2022), allowing one to estimate how much of the neural activity is related to the stimulus and how much is not. However, while this approach might work well to estimate the neural activity related to the *acoustic* processing of natural speech, it is not well-suited for modeling neural responses that reflect the *linguistic* processing of that speech. This is because repeating the same speech stimulus is sure to lead to differences in language-related brain activity due to changes in attention and substantial reductions in linguistic information and predictability across repetitions (Kutas & Federmeier, 2011; Renoult et al., 2012). An alternative strategy is to assume that the brains of different people who speak the same language should produce similar responses – both acoustic and linguistic – to the same stimuli (Hasson et al., 2004; Schrimpf et al., 2021). Indeed, using this strategy, previous studies have estimated the explainable variance in electrocorticography (ECoG) and functional magnetic resonance imaging (fMRI) responses to speech by calculating how accurately a selected (target) subject’s brain signal can be modeled using the signals from other (source) subjects (Schrimpf et al., 2021). Specifically, these studies: 1) fit decoders for a target’s brain signals using different numbers of source subjects; 2) fit curves to model the relationship between the number of source subjects and the target subject’s prediction accuracy; and then 3) extrapolated from those curves to estimate the target subject’s brain activity as if there were an infinite number of source subjects.

In the present study, we tailored this framework for the unique characteristics of scalp EEG. These characteristics include high correlation between neighboring channels (i.e., low spatial resolution); low signal-to-noise ratio^1^; and significant inter-individual differences in the spatiotemporal profiles of responses to sensory stimuli (and speech stimuli in particular; (Di Liberto & Lalor, 2017)). All of these factors complicate the estimation of the explainable variance (or noise ceiling) of EEG on individual channels using correlation and regression based on the same channels in other subjects. As such, our approach involves predicting individual channels of the target subject’s EEG after combining EEG across channels and time lags from the source subjects using methods based on canonical correlation (please see methods for details).

We applied the proposed framework to an EEG dataset collected in 19 native English speakers who all listened to an audiobook version of a classic work of fiction. We found that we could successfully model EEG responses in individual subjects based on EEG from other subjects. Moreover, the accuracy of these inter-subject models increased with increasing numbers of source subjects. Extrapolating this increase allowed us to estimate the noise ceiling in each individual target subject. Finally, we then compared the estimated noise ceiling with the prediction accuracy of a speech features-to-EEG model. Specifically, we fit and tested temporal response functions (TRFs; Crosse et al., 2016) based on the speech envelope, impulses at word onset, and the lexical surprisal of each word. The comparison showed that the estimated noise ceiling was significantly (although not substantially) higher than the speech-to-EEG prediction accuracy. This suggests that the chosen speech feature predictors could explain a large proportion – but, importantly, not all – of the explainable variance in EEG responses to natural speech.

## Methods

### Dataset

To test our approach, we used a publicly available dataset of EEG responses to natural speech (Broderick et al., 2019). These EEG data were collected using a BioSemi ActiveTwo system and included 128 channels of scalp EEG plus 2 mastoid channels sampled at 512 Hz. The EEG was collected from 19 healthy adult native speakers of English who listened to an audiobook version of a classic work of literature read by a single male American speaker of English. Subjects listened to 20 segments of the story each lasting approximately 180 seconds, with each segment continuing the story from where the previous segment ended (i.e., the speech segments were not repeated within subjects). The dataset was preprocessed by band-pass filtering between 0.5 and 8 Hz, downsampling to 64 Hz, and interpolating noisy channels using procedures described in detail in (Dou et al., 2025).

### Estimating the explainable variance in EEG responses to natural speech using inter-subject modeling

The core assumption at the heart of our approach is that brain activity in response to natural speech in one (target) individual can be well modeled based on the brain activity of other (source) people listening to the same speech. We specifically wished to model the target subject’s EEG on each individual channel. Having a noise ceiling estimate for each channel would facilitate the direct comparison of EEG encoding model performance against that ceiling as well as the normalization of those encoding model performances for comparison across subjects.

Perhaps the simplest approach one could conceive of for modeling one target person’s EEG based on several other source people would be to compute a channel- by-channel correlation between the target EEG and the EEG averaged across those source people. However, such a model would surely be suboptimal for two reasons. First, while EEG responses to stimuli are distributed across the scalp in a way that is generally fairly consistent across people (Luck, 2014), every person’s EEG also shows subtle differences in those spatial distributions based on the unique shape of their head and their unique “fingerprint” of cortical organization and folding. And second, while EEG responses to stimuli are also generally somewhat consistent in terms of their temporal profile (Luck, 2014) – as reflected by the existence and timing of many classic event-related potential components – there are also subtle inter-subject differences in that timing and the relative amplitude of different response components. For these reasons, averaging EEG on a particular channel across subjects would surely smear those inter-subject spatial and temporal differences and lead to a generic, suboptimal estimate for the EEG on that channel in the target individual.

An alternative inter-subject modeling approach would be to fit a model for the data on each EEG channel for the target subject based on the data on *all* the channels of the source subjects – which, in our case would be a (# of source subjects x 128 channels)-to-1 mapping. While this could work in principle, it runs a very high risk of overfitting with limited EEG, and it would be very computationally intensive.

With these considerations in mind, we aimed to model the target subject’s EEG using an approach that could allow for spatial and temporal variability in responses across source subjects but would be less likely to result in overfitting and would be computationally tractable. Our overall strategy for doing this is shown in Figure 1. The approach involved first reducing the dimensionality of the 128-channel EEG to just 3 components for each subject. To do this, we used an approach known as multi-way canonical correlation analysis (mCCA; see below for details). Then we fit a linear model – specifically a temporal response function (Crosse et al., 2016; please see details below) – to predict the target subject’s EEG on each channel from the source subjects’ components – which is a (# of source subjects x 3 components)-to-1 mapping. Using TRFs in this way allowed us to account for differences in the temporal profile of EEG responses across subjects. Finally, because our dataset only had 19 subjects – and because of the relatively low SNR of EEG – it was unreasonable to assume that modeling 1 target subject using 18 source subjects would be a good estimate of the noise ceiling. Because of this, we carried out the above inter-subject modeling approach for different numbers of source subjects. The idea was that, as we increase the number of source subjects, the model for the target subject should improve. We could then extrapolate from that pattern to estimate how well we would be able to predict the target subject’s EEG if we had infinite source subjects – following a nice approach used by (Schrimpf et al., 2021) for fMRI.

**Figure 1.**
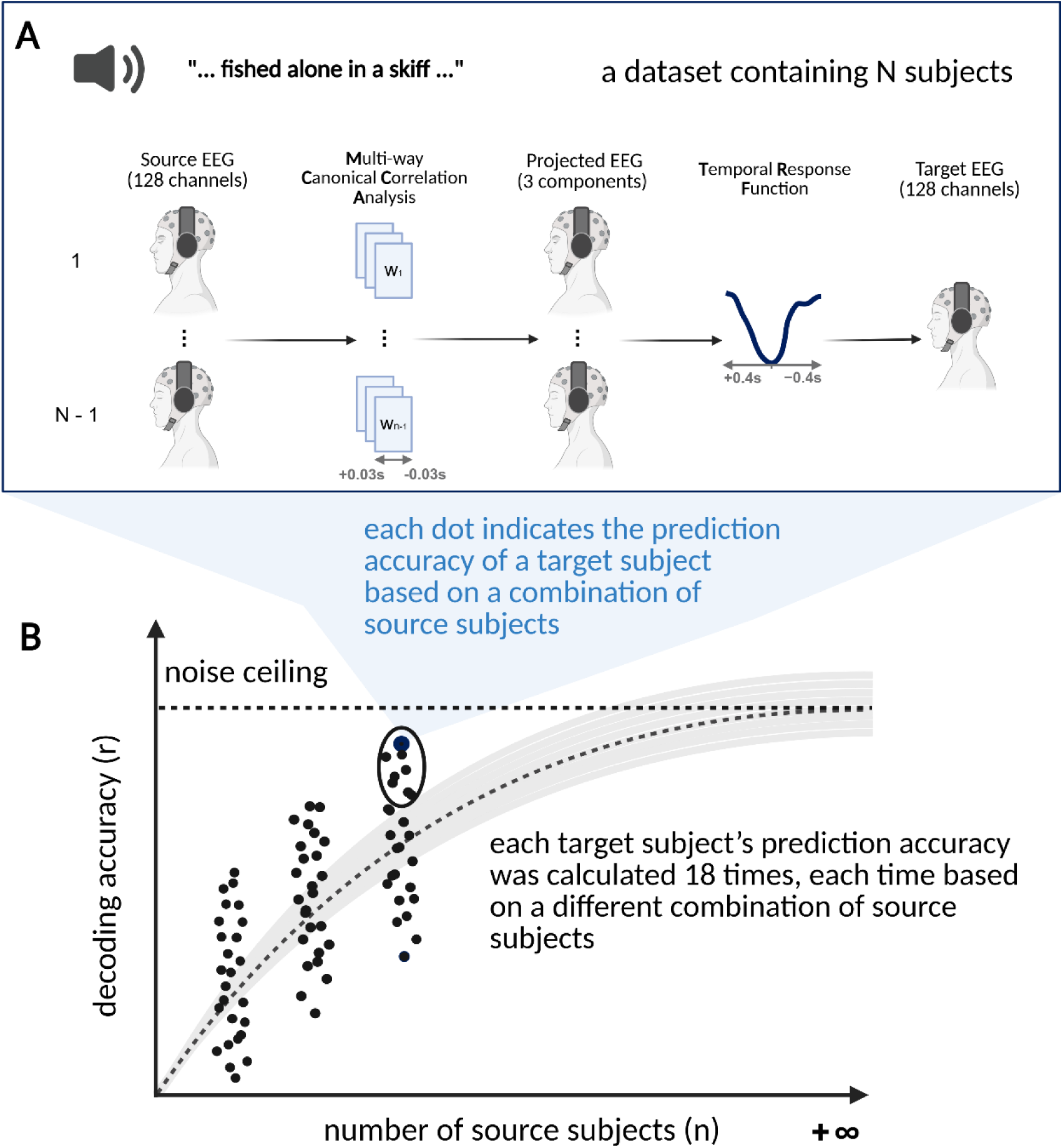
Estimating the explainable variance in EEG responses to natural speech using inter-subject modeling. **A** 19 native speakers of English listened to the same audiobook while their (128-channel) EEG was recorded. One of these subjects was chosen as the target subject. A subset of the other subjects were source subjects (in the figure we show a case with all the other subjects as the source subjects). Multi-way canonical correlation analysis was used to reduce the 128 channel EEG to 3 components for each source subject that were maximally correlated with those of all the other source subjects. A temporal response function was then fit that aimed to model each channel (of 128) in the target subject based on the 3 components in the source subjects (i.e., 128 separate mappings, each with the dimensions: 3 components * # of source subjects-to-1). **B** By increasing the number of source subjects used to model the EEG of the target subject, one can get improved estimates of that target subject’s EEG. Each dot represents the prediction accuracy of the target subject’s EEG based on a different number and specific set of source subjects. The dashed curve represents an exponential function fit to these dots. This exponential curve can be extrapolated to estimate how well the target subject’s EEG could be modeled with infinite source subjects. We take this estimate to be the explainable variance in the target subject’s EEG. Created in https://BioRender.com.

### Accounting for differences in the spatial distribution of EEG across subjects using multi-way canonical correlation analysis

As mentioned above, modeling the data on one EEG channel in a target subject using data from the same channel in source subjects is guaranteed to be suboptimal because the spatial distribution of EEG responses to speech is not identical across subjects. Also as mentioned above, fitting a linear model to map from 128 EEG channels in the source subjects to one channel in the target subject is a many to one mapping that is highly likely to overfit with limited data, as well as being computationally intensive. As such, we wished to account for differences in the spatial distribution of the EEG across source subjects and to reduce the dimensionality of their data before predicting the target subject’s EEG. To do this, we used multiway canonical correlation analysis (MCCA). Specifically, for every specific subset of source subjects, we performed MCCA across that subset of subjects to reduce the dimensionality of their data, and then used the resulting components to predict the data in the target subject.

In brief, MCCA is a method that maximizes the correlation between multiple sets of data matrices through linear projection, and can be solved using two rounds of principal component analysis (PCA; de Cheveigné et al., 2019; Parra, 2018). In the first round, the data matrix of each dataset is whitened using PCA. And in the second round, the whitened principal components of each dataset are concatenated along the component dimension to be jointly decomposed by another PCA. The result is a set of components that are ordered by how highly correlated they are across subjects. In other words, the first few components are maximally correlated across subjects. By choosing a limited number of these, one can greatly reduce the dimensionality of one’s data and retain common signal (i.e., in this case, signal related to processing the speech stimulus) across subjects. Please refer to (de Cheveigné et al., 2019; Parra, 2018) for a detailed exposition of this approach.

To account for differences in the latency of the EEG responses to speech between source subjects, we included time shifts in the implementation of MCCA. This approach allows the algorithm to identify components that are maximally correlated between subjects while allowing for some differences in the absolute latency of those components between subjects. The time shifts were incorporated into the analysis in the same way as has been done for previous MEG responses to speech (Huizeling et al., 2022). Specifically, to balance the computational cost with the ability to accurately model target subjects, we chose a range of time shifts for the MCCA to be -2 to 2 samples (which at a sampling rate of 64 Hz equates to – 31.25 to 31.25 ms). As mentioned before, we chose to focus on the top 3 canonical components for each subject. The reason for using 3 canonical components is that the decoding performance kept increasing as we increased the number of canonical components (CCs) from 1 to 3, but then the performance dramatically dropped with 4 CCs for each subject.

[As an aside, in addition to MCCA, we also tried using principal component analysis (PCA) to reduce the dimensionality of the EEG for each source subject. Specifically, we selected the top 16 principal components for each subject and used those to predict the data on each channel of the target subject’s EEG. However, this procedure did not produce significantly better performance than the MCCA approach and was more computationally expensive (with 16 PCs vs 3 MCCA components; Fig S1).]

### Accounting for differences in the temporal morphology of EEG responses to speech across subjects using temporal response functions

To predict each channel of the target subject’s EEG based on the source subjects’ MCCA components, we used linear (ridge) regression to fit a filter/model – sometimes known as a temporal response function (TRF; Crosse et al., 2016). This approach learns how to weight each MCCA component – at different relative time lags – to predict the EEG for the target subjects with least squares error. Specifically, the TRF is a type of linear time-invariant convolutional kernel that can be used to model how a data point in the output at time *t*_0_ can be predicted by a linear combination of data points in the input within the time range from *t*_0_ + *τ_min_*to *t*_0_ + *τ_max_*, with *τ_min_* and *τ_max_*being the lower and upper bound of the time range. In this study, we aimed to predict the target subject’s EEG (*̂r*) by convolving the TRF weights (ℎ) with the MCCA inputs (*ss*):

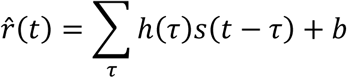

where *τ* indicates the time lag, *b* represents the interception term, and *s* ∈ ℝ*^T^*^,*M*×*K*^with *T*, *M*, and *K* being the length of the stimulus, number of canonical components and the number of source subjects, respectively. To allow for differences in the temporal lag and morphology of EEG responses between the source and target subjects, we set the lower and upper bound of our time lag range to be -400ms and 400ms.

The weights of the TRF can be solved using ridge regression by regressing the output against the time-lagged model input using the following equation:

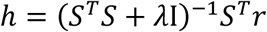

where *S* is the time-lagged version of the model input *s* (i.e., the MCCA components from the source subjects), and *r* is the output (i.e., the EEG responses in the target subject). I is the identity matrix and *λ* is a regularization hyperparameter that is used to control for overfitting and is optimized through cross validation (please see below for details).

As mentioned, the goal of the TRF mapping here was to learn how to weight each source subject’s MCCA components – at different relative time lags – to predict the EEG on each channel in the target subject. By incorporating a range of time lags, we hoped to be able to model the target subject’s EEG responses to speech – which may have their own an idiosyncratic temporal profile – using EEG components from other subjects whose responses might be quite different in terms of their temporal morphology. As such, to confirm that the TRF can be used to map between EEG responses evoked by the same speech stimuli, but that have markedly different temporal profiles, we first did an analysis based on simulated EEG data before applying our approach to the real data in this study (please see results below).

For our simulation, 8 sets of one-dimensional EEG responses were simulated by convolving 8 simulated TRF kernels with a 15-min long acoustic envelope. The range of time-lags for the TRFs was set to -100 – 400 ms. Each TRF kernel was simulated by summing three partially overlapping Gaussian functions that were spread along the time lag axis. The time location of the peak of each of the three Gaussian functions was chosen randomly from two candidate time points, leading to 2^3^ = 8 combinations of locations of the Gaussian functions. Specifically, the two candidate time points for the left, middle and right Gaussian function were 62.5 or 93.8ms, 109.4 or 156.3ms, and 171.9 or 234.4ms, respectively. The width (sigma) of each Gaussian function was selected from a uniform distribution between 1.5 and 3.0. The amplitude values of the left and right Gaussian functions were selected from a uniform distribution between 1.0 and 3.0. And the middle Gaussian function’s amplitude was sampled from -3.0 to -1.0 uniformly. The unbiased accuracy of decoding one simulated EEG timeseries from another simulated EEG timeseries was obtained through the nested cross-validation framework introduced below.

### Estimating the explainable variance in the target subject using different numbers of source subjects

Following the method used in (Schrimpf et al., 2021), we sought to predict a target subjects data using data from different numbers of source subjects, to then fit curves to those predictions as a function of increasing source subject numbers, and to then extrapolate from those curves to estimate the target subject’s brain activity as if there were an infinite number of source subjects.

Predicting the target subject based on all other 18 subjects is straightforward – one just predicts the target subject’s data based on all other available data. Furthermore, when predicting the target subject based on data from a single source subject, there are 18 ways to choose which source subject. However, when predicting a target subject with a number of source subjects between 1 and 18, there is a question of how to choose those source subjects. In this situation, rather than basing our estimates on single subsets of source subjects, we sampled different sets of source subjects. Specifically, given a number of source subjects, *n*, and a selected target subject, *j*, we sampled 18 combinations of *n* source subjects from the remaining 18 subjects in the dataset. (We chose 18 combinations to be consistent with the fact that there were 18 ways to choose 1 source subject.) And, as mentioned, when *n* equals 18, there was only one possible combination. Thus, we obtained a total of 342 (18 * 19, when the source subject number *n* < 18) plus 19 (1 * 19, when the source subject number *n* = 18) sets of prediction accuracies for each EEG channel in each target subject (see Fig 6A below and associated figure caption). Importantly, we generated the predictions of each target subject’s EEG in an unbiased way using a 10-fold nested cross validation procedure (CV; please see details below).

Having generating these EEG predictions across these different samples of different numbers of source subjects, we again followed (Schrimpf et al., 2021) by fitting the following curve to those predictions accuracies using the curve_fit function implemented in SciPy (Virtanen et al., 2020):

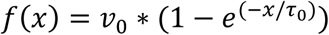

where *v*_0_ and *τ*_0_ are parameters to be fitted, with the lower and upper bounds of *v*_0_ being [0, 1] (i.e., 0% variance explained and 100% variance explained) and the bounds of *τ*_0_ being negative and positive infinity. The fitted value of *v*_0_ for each target subject was taken as the estimate of the total explainable variance in their EEG.

In our initial attempts to fit these curves, we noticed that for many EEG channels that were poorly predicted (i.e., that carried relatively little speech-related activity), the fits of the two parameters described above could be very noisy. In particular, when those two parameters were unconstrained, the *v*_0_ parameter could reach implausibly high values based on extrapolating from very small noisy EEG predictions. To circumvent this problem, we made the assumption that every channel within the same subject should have a similar shaped curve. In practice, this meant that we forced all channels within each subject to share the same *τ*_0_ . In other words, we instead fitted the following function:

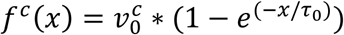

where *c* indicates the *c*th channel. And the parameters for all channels are fitted together by minimizing the following function:

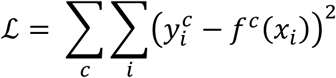

where *y_i_* indicates the *i*th sampled EEG prediction accuracy at the channel *c* and *x_i_* indicates the corresponding number of source subjects.

To estimate the confidence interval of the fitted curves, we implemented bootstrapping to resample pairs of source subject numbers and EEG prediction accuracies, and fit curves to those resampled pairs (Schrimpf et al., 2021). To account for the relatively large inter-individual differences in EEG SNR, we implemented the bootstrapping in a hierarchical manner (Saravanan et al., 2020). In detail, for each round of bootstrapping and for each source subject number *n*: 1) we first resampled the target subjects to include in the curve fitting for current iteration, 2) we then resampled the combinations of source subjects for each resampled target subject, 3) we finally obtained a single prediction accuracy value by averaging across all the sampled prediction accuracies. All resampling was performed with replacement.

### Estimating decoding accuracy unbiasedly using nested cross-validation

As mentioned above, to get an unbiased estimate of the target EEG prediction accuracy, we incorporated a 10-fold nested cross-validation (CV) to fit and test the proposed model. The nested CV consists of an outer CV and an inner CV. Within each of the 10 folds of the outer CV, a 9-fold inner CV was performed to estimate the optimal model regularization. Specifically, in the outer CV, the dataset was divided into 10 splits. Within each fold of the outer CV, 1 split was chosen as the testing data, and the other 9 splits were first applied with the inner CV to find the optimal *λ* and were then used as training data to fit a model with the optimal *λ*. Within each round of the inner CV, 8 of 9 splits were chosen as inner CV’s training data to fit candidate models for each combination of the selected *λ* and the remaining 1 split of data was used to test the candidate models’ prediction accuracies. Thus, when only one hyperparameter needs to be optimized, the model accuracies obtained during an inner CV can be constructed as a two-dimensional matrix with each row representing a proposed hyperparameter and each column representing a fold of the inner CV. The optimal *λ* corresponds to the row that has the largest average-across-column value. The inner CV is completed when all proposed hyperparameters have been used to fit a candidate model and all splits have been used to test the candidate model. The outer CV stops when all the splits have been chosen as the test data. However, to estimate an explainable variance for each channel, a separate best *λ* was identified for each output channel.

### Benchmark Predictors

The primary goal of our study was to establish a framework for estimating the explainable variance of EEG responses to natural speech. However, we also wished to determine how well current speech features-to-EEG models do at explaining that explainable variance. To this end, our estimated noise ceiling was compared with the EEG prediction accuracy of a speech-to-EEG TRF, which encodes EEG signals from pre-selected speech features. The speech features we chose to include were the acoustic envelope, impulses at the start of each word that reflected the lexical surprisal of that word relative to its preceding context, and unit height impulses at word onset.

#### Acoustic envelope

The broadband amplitude envelope was estimated using the Hilbert transform (Di Liberto et al., 2015). It was used to quantify variance in the EEG that is related to acoustic processing.

#### Lexical surprisal impulses

Lexical surprisal is a measurement of how unexpected a word is given its prior words and was calculated as the negative logarithm of the word’s contextual probability. The surprisal values were estimated using a GPT-2 model (Radford et al., 2019), based on considering a maximum of 1024 previous words as context. A detailed description can be found in (Dou et al., 2025). To create the EEG predictor, impulses were placed at the start of each word with a height that corresponded to the lexical surprisal of that word.

#### Word onset impulses

Unit height impulses at word onset were also included as an EEG predictor. They aimed to capture additional acoustic or linguistic variance in the EEG that might be time-locked to the start of words. This could be due to acoustic arise at word level and cannot be explained by lexical surprisal impulses. It was built by placing unit impulses in a zero vector at the time location that corresponds to word onsets.

When predicting the EEG data for each subject using these speech features, we again used 10-fold nested cross validation to optimize the regularization. To be consistent with the approach adopted in many recent studies, we identified a single common *λ* for all channels within each subject.

### Hardware acceleration of computation

To accelerate computing. We reimplemented the MCCA function of the NoiseTool package (de Cheveigné et al., 2018) and mTRFpy (Bialas et al., 2023) using CuPy (Okuta et al., 2017) to run the proposed model on Nvidia’s GPUs.

## Results

### The TRF can account for inter-individual differences in the temporal morphology of brain responses to speech

To first validate whether the TRF can account for variabilities in the temporal response profiles of EEG across subjects, we use the TRF to map between simulated EEG timeseries that were generated under different temporal response profiles. In detail, we set up two types of TRF instances: TRF generators and TRF decoders. The TRF generators were used to generate simulated EEG timeseries. And then the TRF decoders were used to decode one simulated EEG timeseries from another simulated EEG timeseries. As mentioned in the methods section, we simulated 8 EEG timeseries, so there are 28 pairs of simulated EEG timeseries, and thus 28 decoding accuracy values of decoding one EEG timeseries from another one (Figure 2). The decoding accuracy was measured using Pearson’s correlation.

**Figure 2.**
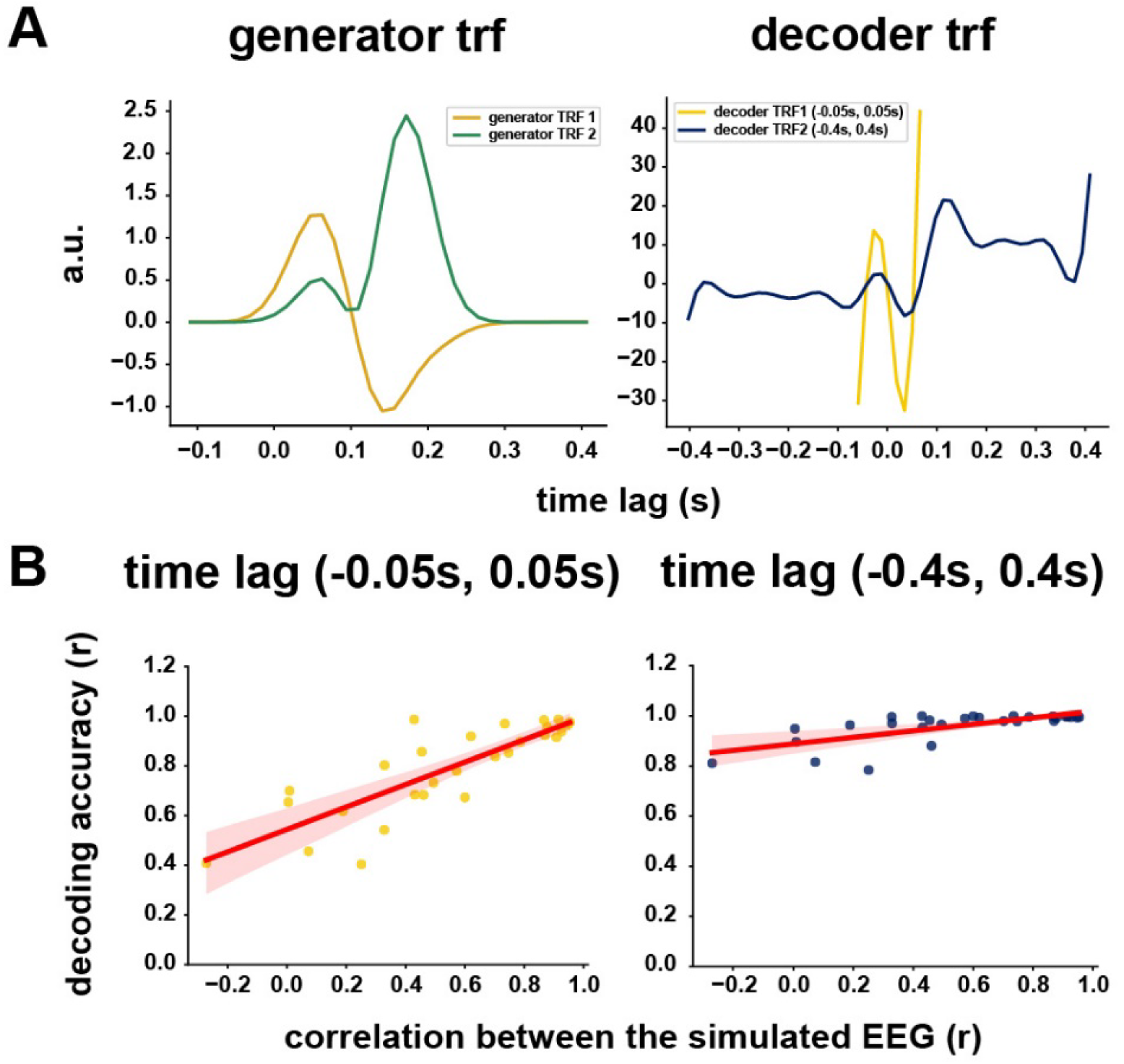
Simulations showing that temporal response functions can be used to model one EEG timeseries based on another EEG timeseries, even when those timeseries were generated based on responses with very different temporal profiles. **A.** Left: An example pair of generator TRFs. The correlation between these generator TRFs was -0.5231. The correlation between the corresponding simulated EEG was -0.2721 (bottom left data point in Fig 2B). Right: The corresponding decoder TRFs with different time-lag ranges. When the time-lag range was -0.05 – 0.05s, the decoder TRF’s prediction accuracy was 0.4082 (bottom left data point in left panel of Fig 2B). When the time-lag range was -0.4 – 0.4s, the decoder TRF’s prediction accuracy was improved to 0.8117 (bottom left data point in right panel of Fig 2B). **B.** Left: When the time-lag range of decoder TRFs was -0.05 – 0.05s, there is a clear trend that the lower the correlation between simulated EEG timeseries the lower the EEG-to-EEG decoding accuracy. Right: When the time-lag range of decoder TRFs was enlarged to -0.4 – 0.4 s, the negative effect of low correlation between simulated EEG on the decoding accuracies was mitigated.

Our simulations specifically sought to explore the question of how well the TRF decoders could do at modeling between two EEG timeseries that were generated using different TRF generators. Unsurprisingly, we found that we could very reliably decode one simulated EEG timeseries from another simulated EEG timeseries when their generative TRFs were very similar. However, when trying to decode one simulated EEG timeseries from another timeseries that was simulated with a very different TRF generator, the accuracy was very low when using a TRF decoder with a narrow range of time lags (−0.05 to 0.05 s; Figure 2B left). However, when using a broader range of time lags (−0.4 – 0.4s; Figure 2B right), we found we could reliably decode one simulated EEG timeseries from another, even when their generative TRFs were very different. For example, when two TRF generators have a very poor correlation with each other (−0.523; Figure 2A left), the correlation between the corresponding pair of simulated EEG timeseries is -0.272 (Figure 2B left, bottom left data point, x-axis). Using a decoder TRF with a narrow range of time lags (−0.05 to 0.05 s) enabled us to decode one of these timeseries from the other with a Pearson correlation of 0.4082 (Figure 2B left, bottom left data point, y-axis). Meanwhile, using a decoder TRF with a broader range of time lags (−0.4 to 0.4 s) greatly improved our ability to decode one of these timeseries from the other (up to a Pearson correlation of 0.8117; Figure 2B right, bottom left data point, y-axis). These simulation results indicate that the TRF can account for variabilities in the temporal response profiles of EEG. Furthermore, it shows that with a longer window of time lags, it is possible to compensate for large differences in temporal response profiles. The averaged decoding accuracy of the TRF with narrow range of time lags was 0.7898, and that of the TRF with broader range of time lags range was 0.9592 (Wilcoxon signed-rank test W = 0.0, p = 7.4506e-09, two-tail).

### Establishing a benchmark by predicting EEG based on speech features

To establish a benchmark against which to compare our estimates of explainable variance, we first estimated the prediction accuracy of EEG responses to natural speech based on certain speech features using a speech-to-EEG TRF. The three chosen features were the acoustic envelope, lexical surprisal impulses, and word onset impulses. We performed 10-fold nested cross-validation to fit and test the model on each subject’s EEG data. The best regularization parameter, *λ*, was selected from 10*^i^*, where *i* ranges from -4 to 3, inclusively. The scalp-level prediction accuracy (Pearson’s r) averaged across both EEG channels and subjects was 0.0507. Prediction accuracy varied across channels consistent with previous studies (Fig 3; Broderick et al., 2018; Di Liberto et al., 2015).

**Figure 3.**
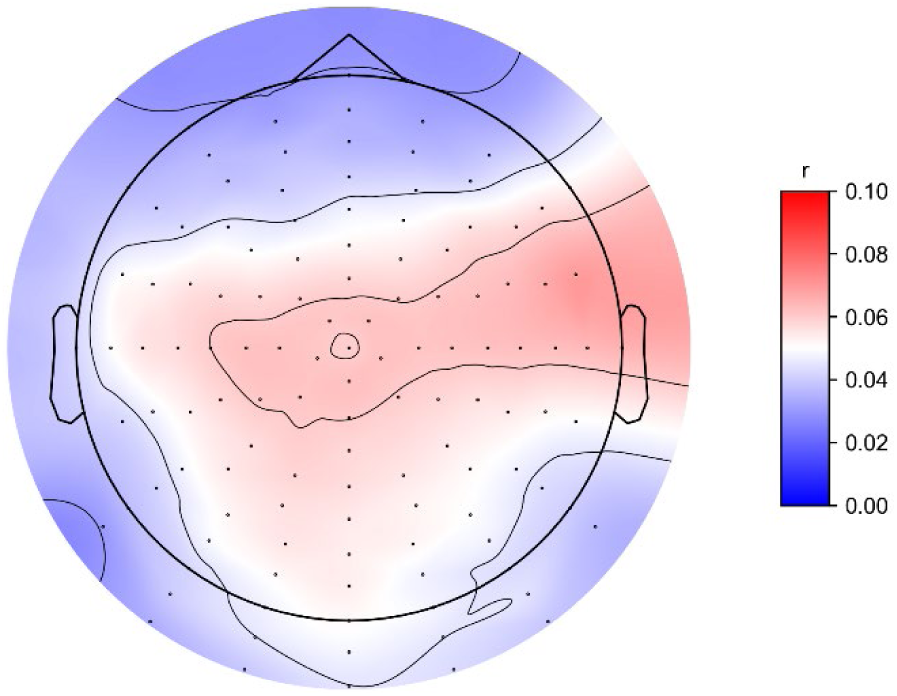
**The prediction accuracy of speech-to-EEG TRF.**

### The estimated noise ceiling is significantly greater than the prediction accuracy based on selected speech features

By modeling the EEG data on each channel in each target subject based on MCCA components from different numbers of source subjects we fit a curve aimed at estimating the total explainable variance in EEG responses to natural speech. We also used a bootstrapping procedure to estimate a confidence interval for this explainable variance. Figure 4 shows the results of this procedure aggregated all subjects and EEG channels. Specifically, before the hierarchical bootstrapping, the sampled EEG-EEG prediction accuracies were averaged across channels for each subject. During the hierarchical bootstrapping, results from all subjects were pooled together for the hierarchical resampling. The estimated median explainable variance (the dotted horizontal line in Figure 4) across all channels and subjects was 0.0608, with a confidence interval of 0.0491 – 0.0796 (based on *α* = 0.05, two-tail). The blue horizontal line indicates the speech-EEG model’s prediction accuracy averaged across subjects.

**Figure 4:**
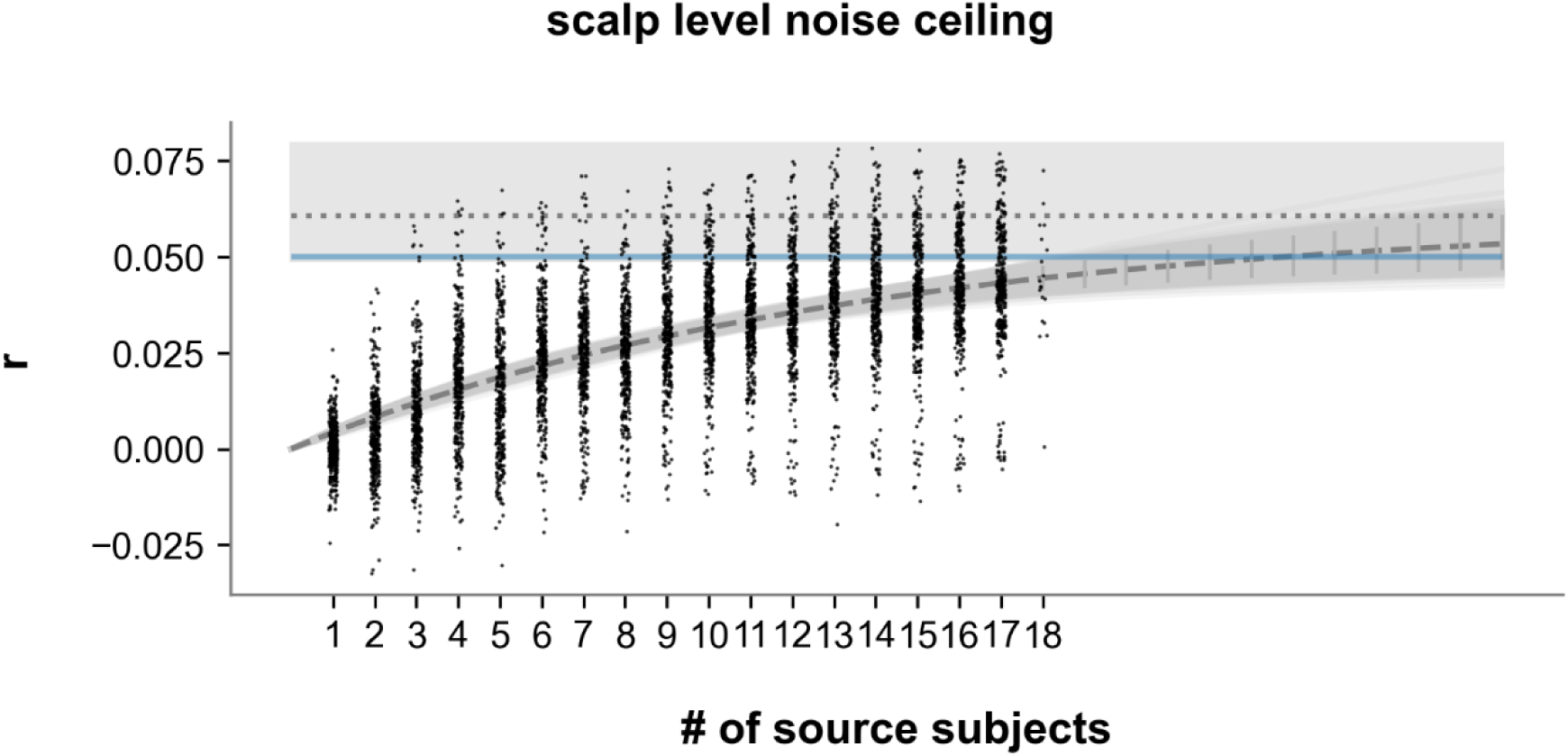
The explainable variance in EEG responses to natural speech aggregated across subjects and channels. Each dot represents the accuracy of decoding one target subject’s EEG from a group of source subjects’ EEG. Grey-colored lines indicate curves fitted during each iteration of hierarchical bootstrapping. The dashed line indicates the median of the fitted curves during hierarchical bootstrapping. The blue horizontal line indicates the speech-EEG model’s prediction accuracy averaged across subjects. The dotted horizontal line indicates the median of the estimated explainable variance aggregated across all subjects and channels.

To estimate the group-level noise ceiling for each individual EEG channel, we applied the same hierarchical bootstrapping procedure to samples of prediction accuracies pooled across all subjects without averaging across channels. The left panel of Figure 5 visualizes the median and the lower bound of the confidence interval of the *v*_0_ parameter that was estimated using the hierarchical bootstrapping. We also compared the noise ceiling with the speech-to-EEG accuracies (Figure 5 right). The white dots indicate the sites where speech-to-EEG accuracy is smaller than the lower bound of the confidence interval of the noise ceiling. The results indicated that the selected features cannot explain all the variance of EEG signals that reflect speech processing. It should be noted that in this group-level case, we consider all subjects together as an average subject, so still a single *τ*_0_ parameter was fitted.

**Figure 5.**
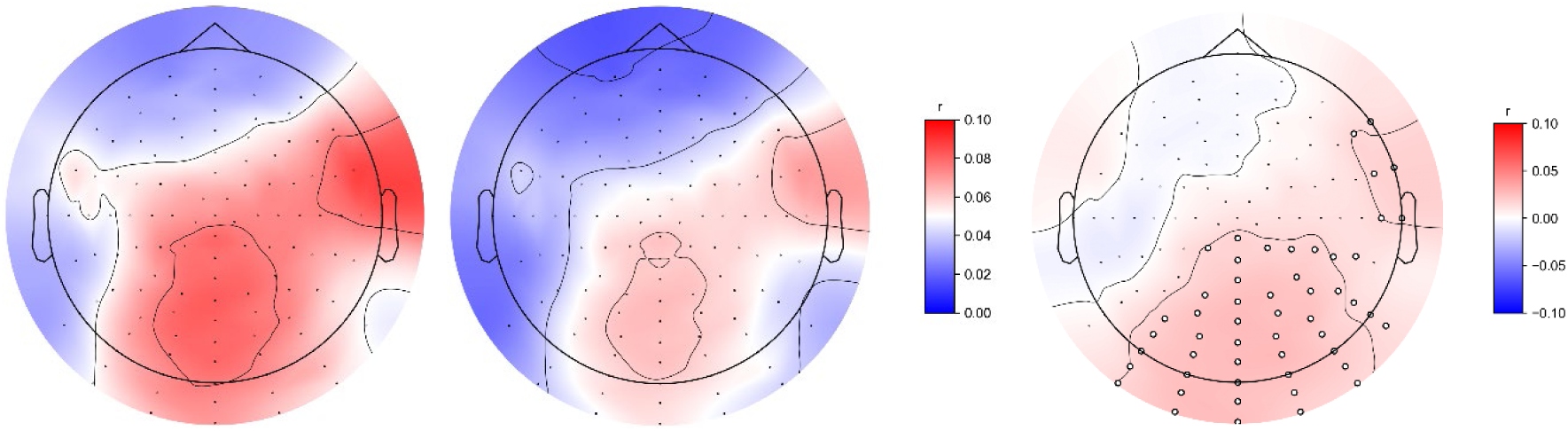
Estimated noise ceiling cannot be fully explained by the speech-to-EEG model. Left: The median of the confidence interval of the group-level channel-wise noise ceiling. Middle: The lower bound of the confidence interval of the group-level channel-wise noise ceiling. Right: The estimated noise ceiling minus the speech-to-EEG prediction accuracy. Channels with significant positive differences are labeled with white dots.

**Figure 6.**
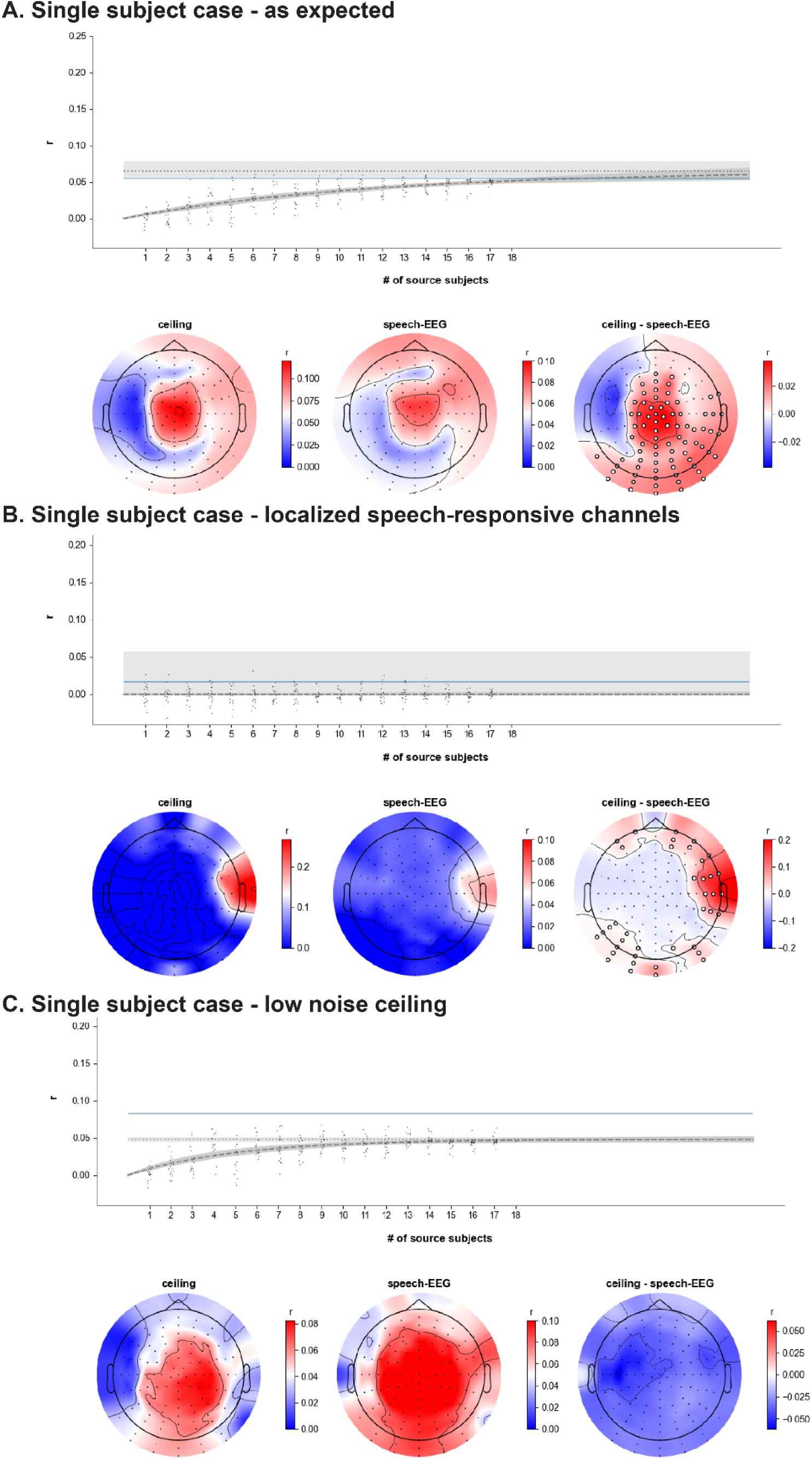
Single-subject level noise ceiling estimations. A. The case when the estimated noise ceiling shows an expected pattern. B. The case of an estimate for a subject having very localized speech EEG responses. C. The case when the estimated noise ceiling is lower than the speech-EEG model’s prediction accuracy.

A central motivation for wanting to estimate the explainable variance in EEG responses to speech is to be able to better interpret inter-subject variability and what constitutes optimal model performance in an individual subject. With that in mind, we also examined the performance of our noise ceiling procedure at a single-subject level. It should be noted that here we used normal bootstrapping because it involved fitting a curve for a single subject. Here, we present three example cases (Figure 6). The first case (Figure 6A) displayed sensible behavior with a median noise ceiling of 0.0648 and a confidence interval of 0.0573 – 0.0776. The speech-to-EEG prediction for this subject was 0.0551. An example case of an estimate for a subject having very localized speech EEG responses is shown in Figure 6B. In this case, the procedure produced a scalp-level noise ceiling estimate with a median of 0.0001 and a confidence interval of 0.0000 – 0.0571. The scalp-level speech-to-EEG prediction for this subject was 0.0125, which is very low and suggests that our approach estimated a zero scalp-level noise ceiling here simply because the EEG responses for this subject were very weak across all channels. However, channel-level speech-EEG has a higher prediction accuracy at the right temporal region, suggesting that the speech response of this subject is very localized. Consistent with this, the similar spatial pattern shown in our noise ceiling estimation indicates that the proposed method still works in the localized case. We also show a case of a low noise ceiling estimate with a median of 0.0477 and a confidence interval of 0.0458 – 0.0504. The speech-to-EEG prediction for this subject was 0.0825. The confidence intervals mentioned above were estimated with the hierarchical bootstrapping method with *α* = 0.05, two-tail. This case indicates that, although the proposed method estimates a group-level channel-wise noise ceiling that is higher than the speech-EEG model’s prediction accuracy, it can still underestimate the noise ceiling in some cases. When testing this method on each subject separately, 11 of 19 subjects were found to have a scalp-level noise ceiling that is significantly higher than the corresponding benchmark. However, at the channel level, almost all subjects have a subset of channels at which the noise ceiling is significantly higher than the corresponding benchmark. Please refer to supplementary table S4 and supplementary figure S5 for detailed scalp- and channel-level noise ceiling results for each subject.

## Discussion

### Motivation, methods, and results

A classic way to estimate the noise ceiling of brain responses to stimuli is through stimulus repetition. The noise ceiling of a subject is calculated as the correlation between brain signals collected from that subject during the first and second presentations of the same stimulus. This method assumes that brain responses stay consistent across stimulus repetitions. However, although this assumption may hold true for low-level sensory stimuli, it is very unlikely to apply to the case of natural speech. Previous studies have shown that word repetition can reduce the amplitude of word-level EEG responses (Kutas & Federmeier, 2011). In addition, both the sensory and linguistic processing of speech are likely to be strongly affected by stimulus repetition because of reduced attention to the stimulus. Previous studies have shown that strong attention effects on EEG indices of both low-level (O’Sullivan et al., 2015; Power et al., 2012); and higher-level (Brodbeck et al., 2018; Broderick et al., 2018; Teoh et al., 2022) processing of speech.

Our study suggests that without repeating stimuli, the explainable variance (noise ceiling) of EEG responses to natural speech can be estimated by decoding a subject’s EEG from a group of other subjects who were presented with the same stimuli. This method is based on the assumption that the brains of different people who speak the same language should produce similar responses to the same stimuli (Hasson et al., 2004; Schrimpf et al., 2021). This assumption is central to other popular methods for indexing brain responses to natural stimuli, such as inter-subject correlation (Hasson et al., 2004; Kumar et al., 2024), which involves calculating the correlation between the brain recordings of a target subject with the averaged brain recordings of other subjects. This method works well for signals like fMRI where inter-individual differences in the hemodynamic response function are relatively small and where there are well validated methods for spatially normalizing brain maps (Avants et al., 2011). However, the inter-individual variability in the spatial and temporal morphology of EEG responses to stimuli (Luck, 2014) limits the effectiveness of this method for estimating the noise ceiling in EEG. As such, we have adapted an approach for quantifying the inter-subject similarity of brain responses to the same stimuli through cross-subject regression (Schrimpf et al., 2021), which 1) fits decoders for a selected subject’s brain signal (target) using different numbers of source subjects; 2) fits curves to model the relationship between the number of source subjects and the target subject’s decoding accuracy. In our adapted approach, we account for inter-individual differences in the spatial pattern of EEG responses by reducing 128 channels of source subjects’ brain signals to 3 canonical components and that mapping from those components to each individual channel of the target subject’s brain signals.

Our approach takes into account inter-individual differences in the temporal domain by including time-lagging in the decoding of a target subject’s EEG. This time-lagged version of ridge regression is sometimes referred to as the temporal response function (TRF). The importance of accounting for temporal inter-individual variability is also supported by our simulation results, which show that increasing the time-lag range of TRF can improve the accuracy of decoding a simulated EEG signal from another EEG signal with a quite different temporal profile.

However, including all the channels of time-lagged EEG in decoder fitting is impractical for datasets with dense EEG channels and a large number of subjects. Take the dataset used in this study as an example: decoding a target EEG from 18 subjects using a time lag range of -0.4s to 0.4s (at a sampling rate of 64Hz) results in an input dimension of 119,808. Such a large input dimension can lead to heavy computational resource consumption and an increased risk of overfitting. To reduce the dimension of data input to the decoder of the target EEG, data compression methods such as principal component analysis (PCA) can be used to summarize the dataset. However, PCA ignores the correlation across multiple data matrices. To better identify EEG responses that are shared across subjects and reduce the data dimension, this study applied multi-way Canonical Correlation Analysis (MCCA) (Arana et al., 2020; de Cheveigné et al., 2019; Huizeling et al., 2022; Parra, 2018) to the source subjects’ EEG to compress the source EEG. To account for differences in the temporal profile of responses between source subjects, we also included a short range of time lags in the MCCA.

To address inter-individual differences in SNR, we used a hierarchical bootstrapping method to estimate the confidence interval of the noise ceiling. One modification this study made on curve fitting is that, based on the assumption that channels within the same subject should have a similar shape of the curve, all channels of the same subject were forced to share the same *τ*_0_ parameter. At the group level, i.e., all subjects’ prediction accuracies were pooled together when fitting curves, we showed that neither the median, nor the lower bound of this group-level noise ceiling is fully explained by a speech-to-EEG model that includes the acoustic envelope, lexical surprisal impulses, and word onset impulses. This is supported by Figure 5, where, at the group level, the noise ceiling cannot be statistically fully explained by our speech-EEG model at centroparietal and right temporal scalp regions. At the individual subject level, however, we noted some variability in how well the approach worked. We presented three examples of estimated noise ceilings: an expected case (Figure 2.6A), a highly localized speech-response case (Figure 2.6B), and a low ceiling case (Figure 2.6C). The expected case shows a noise ceiling with a spatial pattern similar to that of the speech-to-EEG model – channels with high prediction accuracy were also estimated with high noise ceiling values. The localized speech-response case showed that an almost flat zero line was fitted at the scalp level, indicating that on average the SNR is too low. However, the estimated noise ceiling at the channel-level suggests the proposed method still works even if the sites reflecting speech responses are very localized. The low case showed a noise ceiling having similar spatial pattern as the corresponding speech-EEG accuracy. However, it is lower than the corresponding speech-EEG accuracy, indicating that the EEG of that subject is too different from those of the other subjects to be well decoded. Noise ceiling results for each subject can be found in Supplementary Table S4 and Figure S5. In general, 11 of 19 subjects were estimated with noise ceilings that are significantly higher than the corresponding benchmark and the curve fitting approach failed outright for 1 of 19 subjects. However, a non-significant scalp-level noise ceiling in any individual subject may simply be because so few EEG channels were well predicted by the source subjects. This was supported by the fact that at channel-level, almost all subjects have a subset of channels whose estimated noise ceilings were higher than the corresponding speech-EEG prediction accuracy.

### Limitations and future directions

Our approach has a number of limitations. In the first instance, this study assumes that listeners generate similar EEG responses to the same stories. However, this assumption may be valid only when other factors are controlled. For example, within and between subject variability in attention, comprehension, and familiarity with the stimulus are all likely to add noise to the process. To test how these factors may affect the similarity of EEG responses across subjects, relevant behavioral measures may be included in future data collection. Additionally, even when all related factors are controlled, this assumption may not hold for some subjects. Given the non-invasive nature of EEG, different subjects will naturally vary in the strength of both their general auditory and speech/language specific responses to natural speech (Prinsloo & Lalor, 2022). And, for subjects whose scalp responses differ greatly from others – or are just very weak – it may be that the approach cannot do a good job of estimating those responses (e.g., the cases in Figure 6B & C). Indeed, the proportion of people who do not produce clear sensory and linguistic EEG responses to natural speech is unknown. This is a question for future work.

The simulation results indicate that increasing the time lag range of the decoder TRF can accommodate larger inter-individual differences in the EEG response profile. However, increasing the time lag range will include more parameters in the decoder, leading to higher computational cost and an increased risk of overfitting. Thus, a future direction for optimizing this method is to search for an optimal time lag range for each decoder between each pair of source and target EEG timeseries. An optimal time lag range also provides meaningful information about the degree of difference between the EEG response profiles of two subjects. Related to this issue, another potential limitation of our study was that – for computational reasons – we downsampled our data to 64 Hz (from an initial sampling rate of 512 Hz). Lowering the sample rate in this way may have made us less sensitive to subtle differences in the temporal morphology of EEG responses across subjects.

The proposed approach relies on MCCA to construct a customized spatiotemporal filter for each source subject to reduce the data matrix’s dimensionality. Although MCCA performs better than PCA at extracting shared signals across subjects, we showed that this method can underestimate the noise ceiling values for a subset of subjects. Additionally, we found that although the decoding performance kept increasing as we increased the number of canonical components (CCs) from 1 to 3, the performance dramatically dropped with 4 CCs for each subject. This may indicate that, given limited source subjects, MCCA tends to be overfit as the number of CCs goes up. A future direction for testing the effectiveness of MCCA is to directly identify which speech features are represented in the CCs by predicting those CCs with speech features.

Another potential limitation of our work is that we fit the MCCA and TRF models as separate steps. An alternative approach would be to jointly optimize the dimension-reduction module and the decoding module using gradient descent and use the prediction accuracy of the target EEG as the objective function. Such an approach might lead to improved estimates of the noise ceiling in individual subjects. In addition, further improvements might derive from including nonlinear activation functions rather than relying on linear models. This might also have the added advantage of obviating the need for using large windows of time lags to account for large inter-subject temporal differences in EEG responses.

Another tunable component of this framework is the curve function for modeling the relationship between the number of source subjects and the decoding accuracy of the target subject’s EEG. In the current study, we used the same natural exponential function as the previous study (Schrimpf et al., 2021). However, the rationale for assuming an exponential relationship between the two variables may need further exploration in the future, as there are other functions that are monotonic and have lower or upper bounds. In this study, we focused on both the median and the lower bound of the estimated noise ceiling. The gap between the median and the speech-to-EEG predictions could be considered the variance in the EEG that remains to be explained by future models. However, given the relatively small number of subjects in our study, that difference should be treated with caution. As such, our statistical analysis focused on comparing the speech-to-EEG predictions with the lower bound of our noise ceiling estimate. This was more conservative but still indicated that the speech-to-EEG model could not explain all of the explainable variance in the data.

Another key limitation of the present study centers on our speech-to-EEG benchmark. Specifically, we have compared our noise ceiling estimates to a speech-to-EEG model based on only the acoustic envelope, lexical surprisal impulses, and word onset impulses. This ignores a wealth of speech features that have previously been shown to be useful in modeling EEG/MEG responses to speech (Brodbeck et al., 2018; Di Liberto et al., 2015; Gwilliams et al., 2025; Teoh et al., 2019). Moreover, our speech-to-EEG model is also based on an assumption that the relationship between the speech features and the EEG is well captured by a linear model. It is clear that brains are not linear, time-invariant systems. Indeed, several studies have shown that EEG is better modeled when relaxing the LTI assumption in such models (Akram et al., 2017; Dou et al., 2025; Keshishian et al., 2020). As such, our claim that current speech-to-EEG models cannot explain all of the explainable variance in EEG responses to natural speech is not a general one. Rather, in this study, we simply show that a linear model based on the envelope, lexical surprisal, and word onset does not even reach the lower bound of our estimate of explainable variance.

Finally, our study focused on estimating the explainable variance in relatively low-frequency EEG data between 0.5 – 8 Hz. However, previous studies have shown that EEG bands above 8 Hz may also reflect speech and language processing. For example, it was found that a decreases in alpha power is related to the difficulty of constructing the syntactic structure of noun-phrase sentences when encoding those sentences into working memory (Vassileiou et al., 2018). Lower alpha power in EEG was also found to be related to the binding of bigrams into phrases (Bastiaansen & Hagoort, 2006). In addition, it was found that integrating lexical-level speech features into unified representations of utterances correlates with changes of power in the beta and gamma bands (Bastiaansen & Hagoort, 2006). Envelope tracking was also found in the gamma band of EEG for some subjects during natural speech comprehension (Synigal et al., 2023). Thus, it is very possible that EEG bands above 8 Hz also reflect speech processing during natural speech comprehension. A future direction would be to test if the proposed method also works on estimating the noise ceiling of speech-processing related EEG responses within bands above 8 Hz.

In sum, we have introduced an approach for estimating the noise ceiling in EEG responses to natural speech – based on the assumption that the best model of the brain responses to speech in one subject is the brain responses in other subjects listening to the same material. By predicting the EEG data in target subjects based on different numbers of source subjects, and then extrapolating to estimate how well we could explain the data with infinite source subjects, we show that the noise ceiling in EEG responses to nature speech is higher than what is predicted by a particular speech-to-EEG model.

## Acknowledgements

This work was supported by a grant from the National Institute of Deafness and Other Communication Disorders (R01 DC021140) and by the Del Monte Institute for Neuroscience. The authors thank Dr. Aaron Nidiffer, Dr. Liberty Hamilton, Dr. Alexander Huth, and Dr. Jonathan Brennan for helpful discussions. We would like to acknowledge technical support from the Center for Integrated Research Computing at the University of Rochester.

## Appendix

**Supplement 1:**
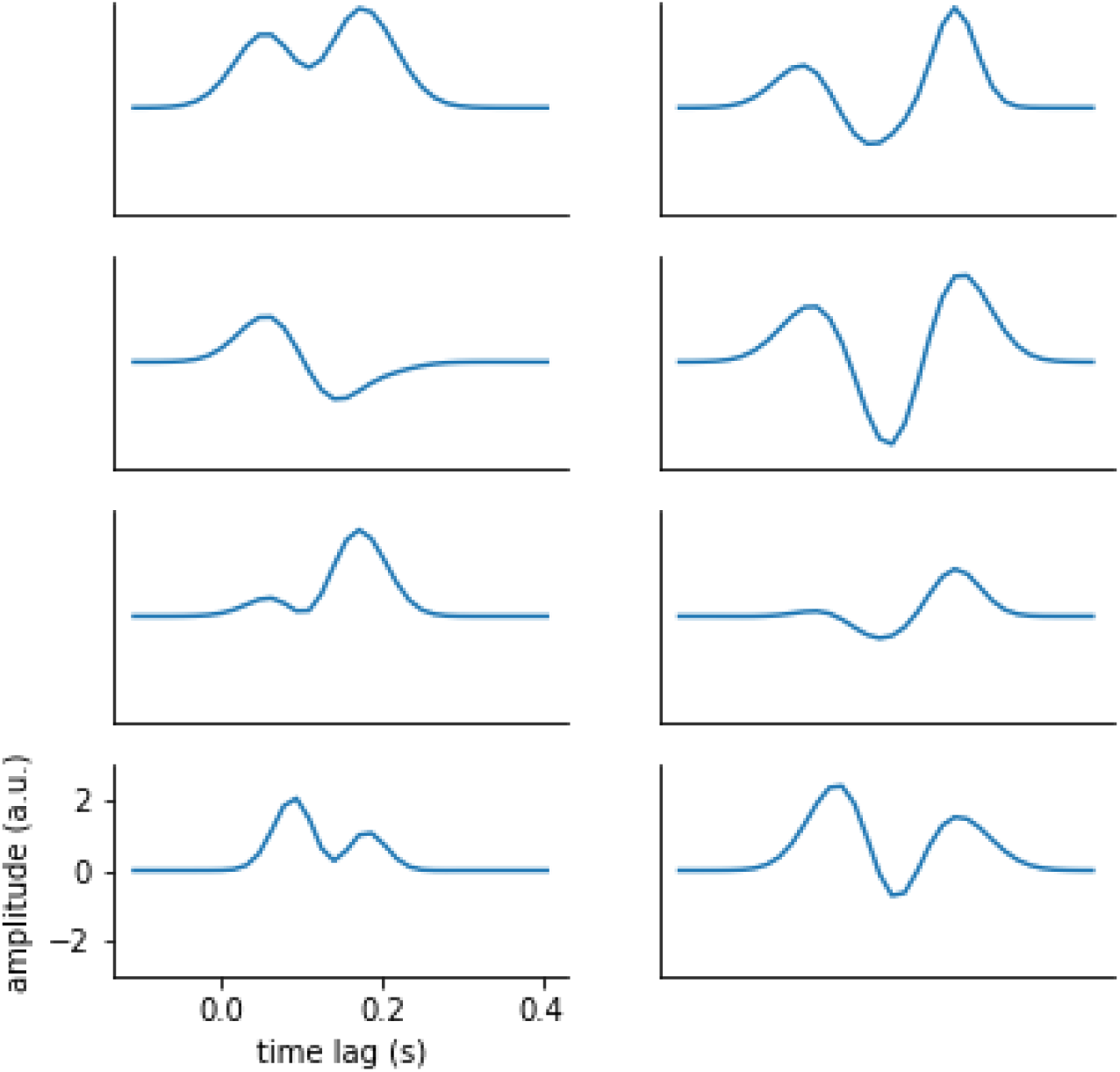
All simulated TRF kernels.

**Supplement 2:**
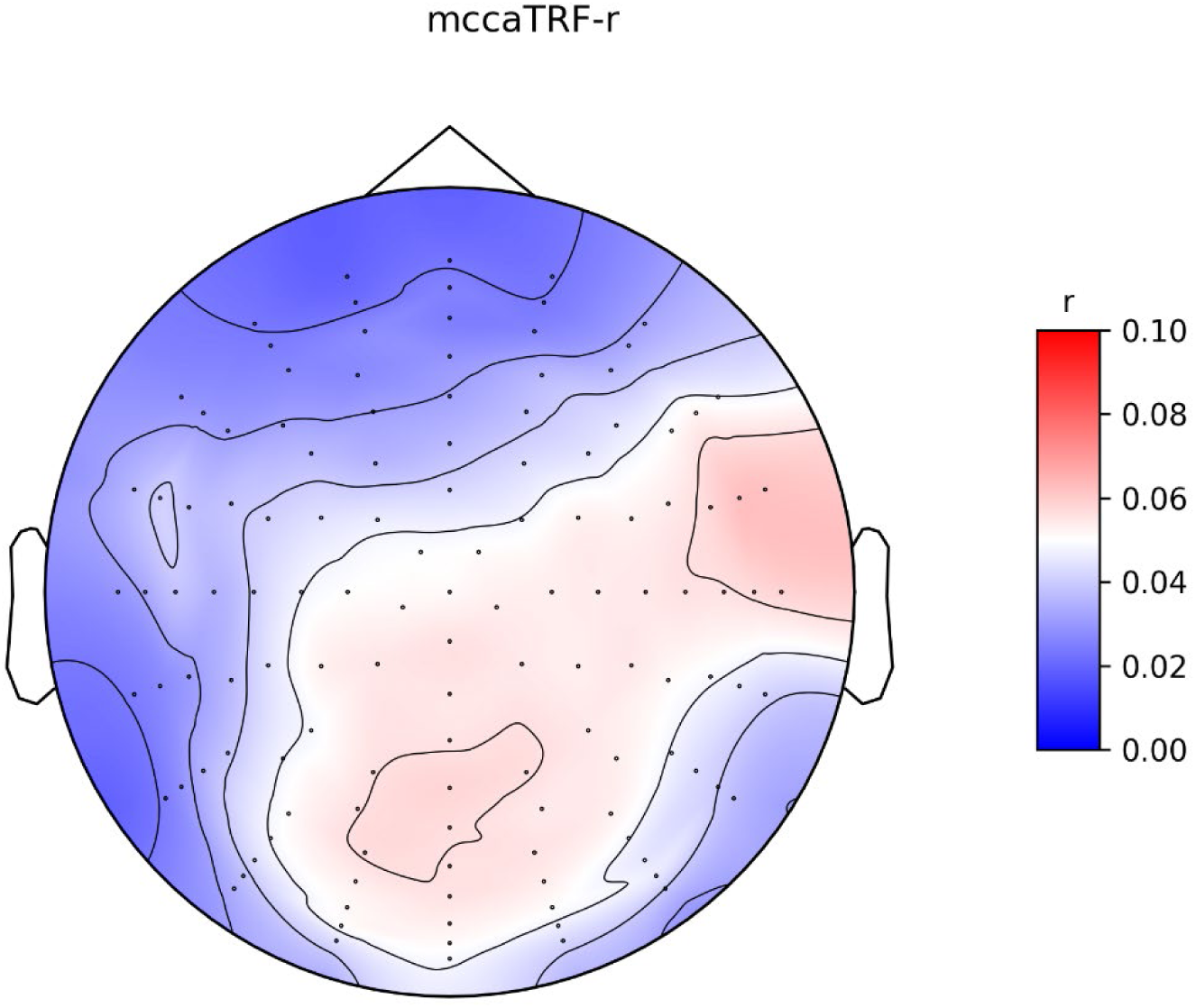
The average decoding accuracy of EEG using all subjects and the MCCA+TRF method.

### PCA is less efficient than MCCA in reducing EEG dimensions

#### Principal Component Analysis

PCA is a method that transform a dataset into a new space where the first “axis” explained the most variance in the dataset, and so on. The benefit of using PCA is that we can choose to use the first N components to keep variance in the original dataset as much as possible while reducing the dimensions of the data. PCA can be solved using eigen decomposition. The covariance of the EEG data can be decomposed into eigen values and eigen vectors, where eigen vectors represent the principal component that represents a basis in the new coordinate and the corresponding eigen values represent the variance explained by this component. In practice, we fitted a separate PCA for each subject within each round of inner CV. During the fitting stage of inner CV, all trials of EEG of one subject within the training splits were concatenated along the time axis before being transformed with PCA that is implemented in the scikit-learn toolbox (Pedregosa et al., 2011). We selected the first 16 principal components based on empirical results.

#### Result

We found that PCA tends to need more components to capture enough information for decoding the target EEG. By comparison, we found that PCA did not perform better than MCCA when using 18 source subjects’ EEG to predict one target subject’s EEG (S3). To balance the fitting time and the information loss, we finally chose 16 components for each subject, with the TRF time-lag range set to -0.05s – 0.05s. We did not notice improvement of performance when using selecting the components explaining the first 99% variance for each subject or when using a separated optimal component number for each subject that was searched using cross-validation. It could be because including more components increases the risk of over-fitting. The statistical test was performed using the Wilcoxon signed-rank test, with p-value corrected by the Benjamini/Hochberg method (*α* = 0.05).

**Supplement 3.**
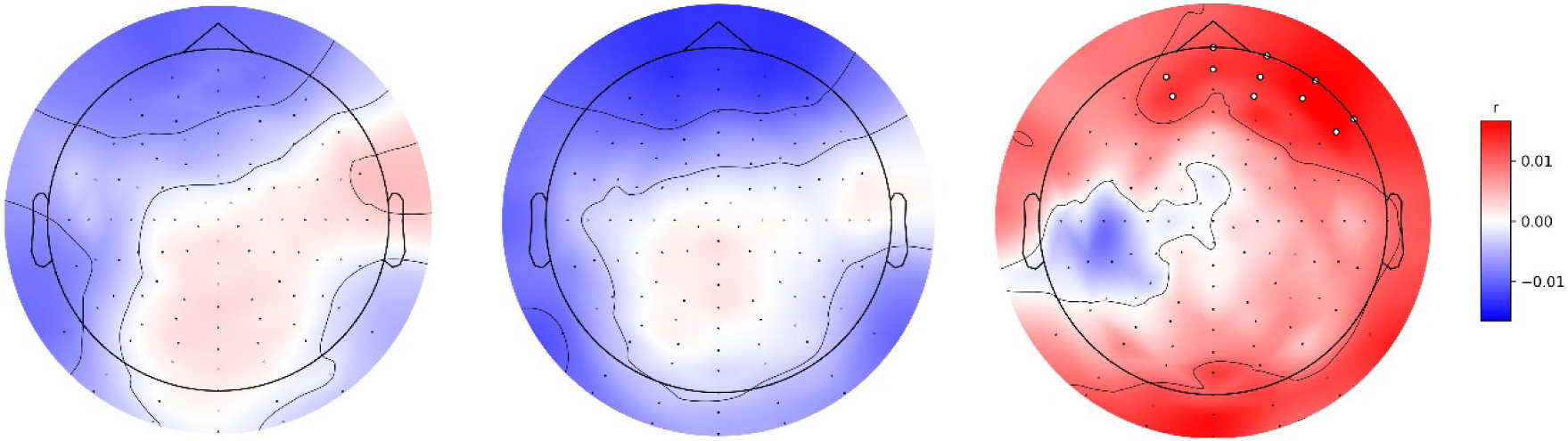
PCA did not perform better than MCCA in decoding target EEG. Left: Group-level TRF decoding accuracy from 18 source subjects when combined with MCCA. Middle: Group-level TRF decoding accuracy from 18 source subjects when combined with PCA. Right: Whited dots indicated the channels showing significant benefits of MCCA. The statistical test was performed using single-side Wilcoxon signed-rank test, with p-value corrected by the Benjamini/Hochberg method (α=0.05).

Summary table of scalp-level for each subject.

Summary table of scalp-level for each subject
| Subject | Speech-EEG | Ceiling | Confidence Interval |
| --- | --- | --- | --- |
| 1 | 0.0478 | 0.0623 | (0.0559, 0.0730) |
| 2 | 0.0291 | 0.0644 | (0.0550, 0.0805) |
| 3 | 0.0432 | 0.2069 | (0.1126, 0.9942) |
| 4 | 0.0551 | 0.0648 | (0.0573, 0.0776) |
| 5 | 0.0631 | 0.0504 | (0.0462, 0.0565) |
| 6 | 0.0506 | 0.0440 | (0.0399, 0.0503) |
| 7 | 0.0369 | 0.0501 | (0.0413, 0.0668) |
| 8 | 0.0491 | 0.0477 | (0.0383, 0.0706) |
| 9 | 0.0443 | 0.1222 | (0.0942, 0.1860) |
| 10 | 0.0706 | 0.0653 | (0.0535, 0.0875) |
| 11 | 0.0165 | 0.0001 | (0.0000, 0.0571) |
| 12 | 0.0825 | 0.0477 | (0.0458, 0.0504) |
| 13 | 0.0841 | 0.0856 | (0.0802, 0.0922) |
| 14 | 0.0514 | 0.0625 | (0.0577, 0.0690) |
| 15 | 0.0420 | 0.0975 | (0.0866, 0.1135) |
| 16 | 0.0584 | 0.0827 | (0.0732, 0.0977) |
| 17 | 0.0399 | 0.0572 | (0.0510, 0.0668) |
| 18 | 0.0333 | 0.0366 | (0.0311, 0.0463) |
| 19 | 0.0524 | 0.0642 | (0.0538, 0.0817) |

**Supplement 4.** Summary of scalp-level speech-EEG prediction accuracy, noise ceiling and confidence interval of noise ceiling.

**Supplement 5.**
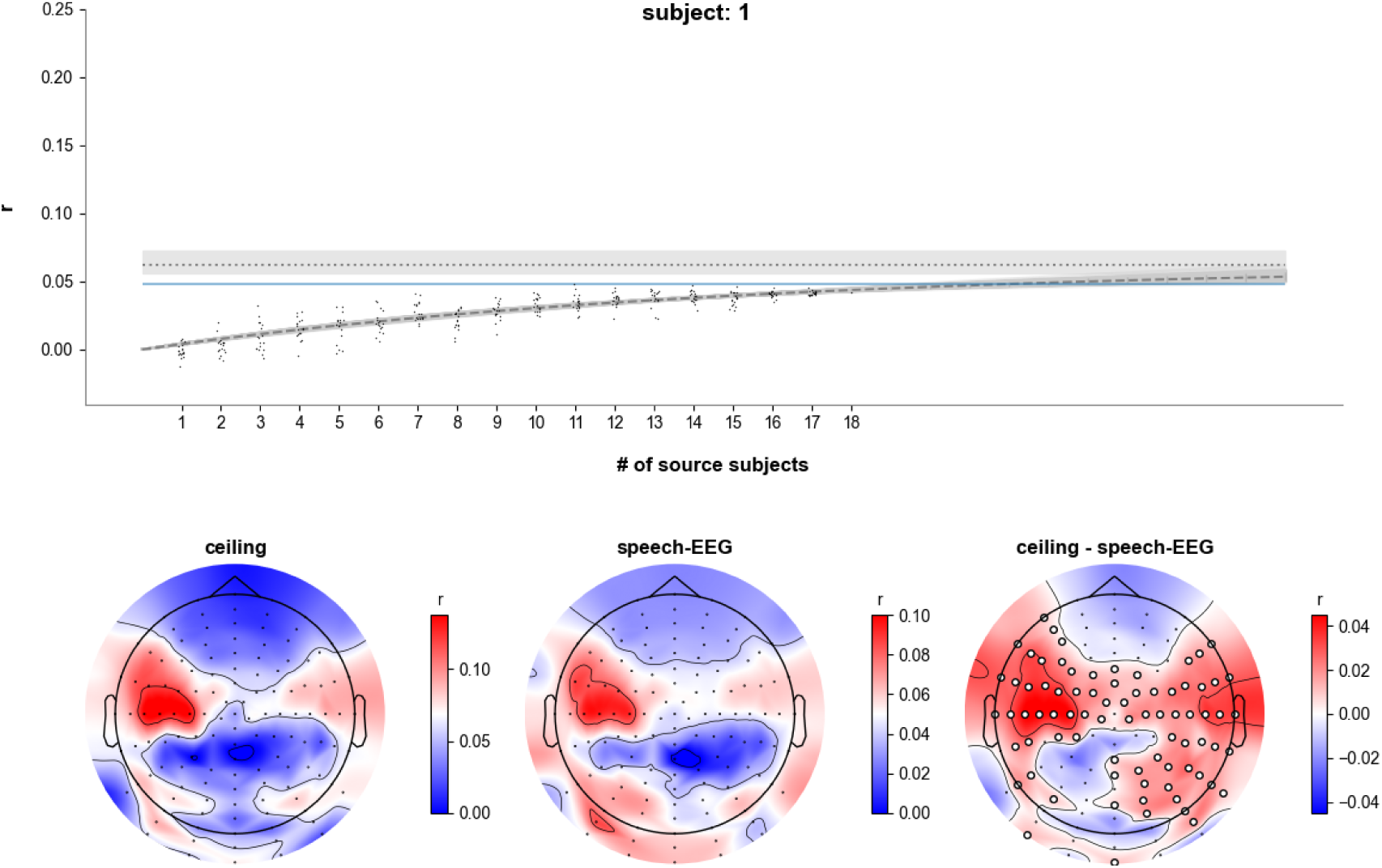
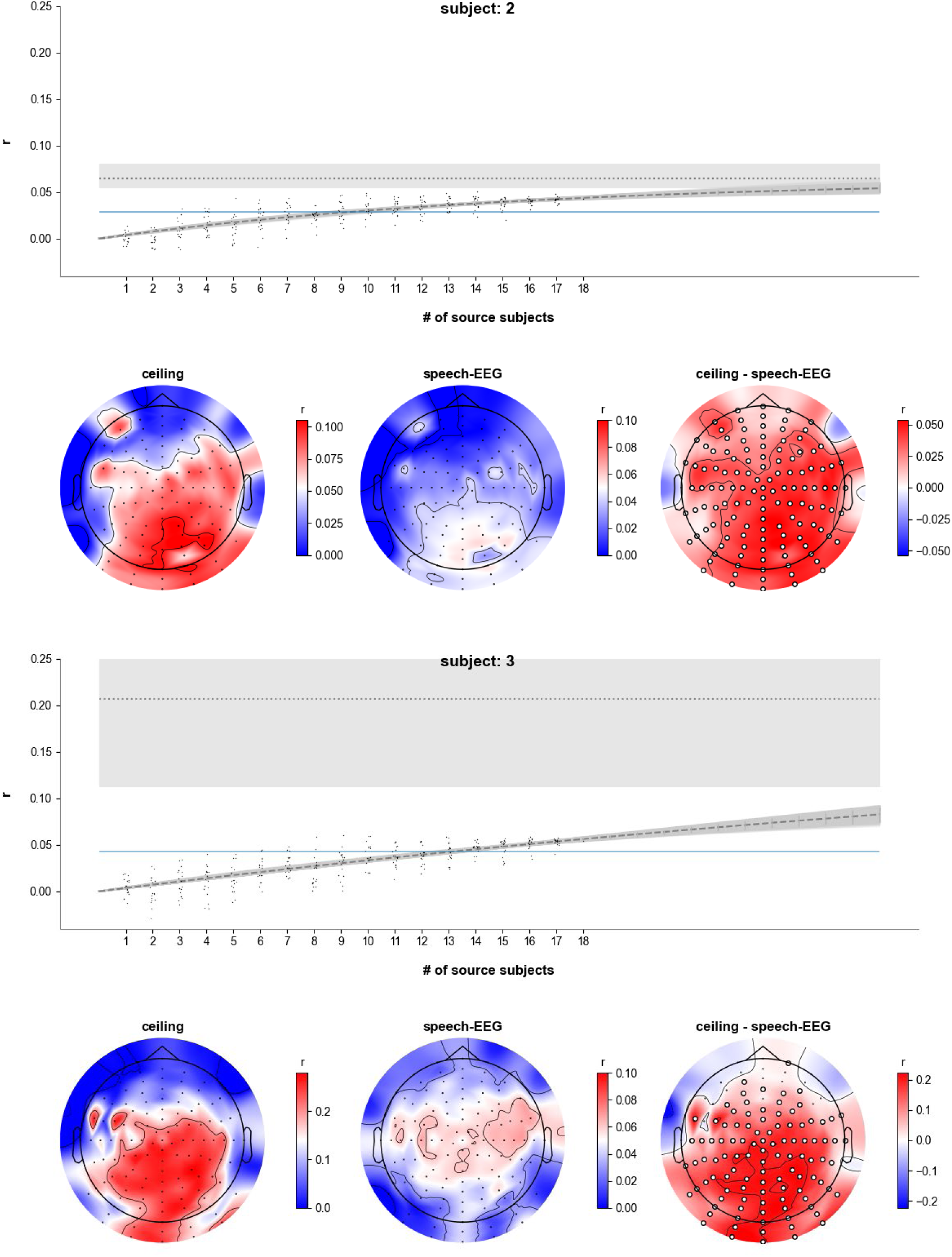
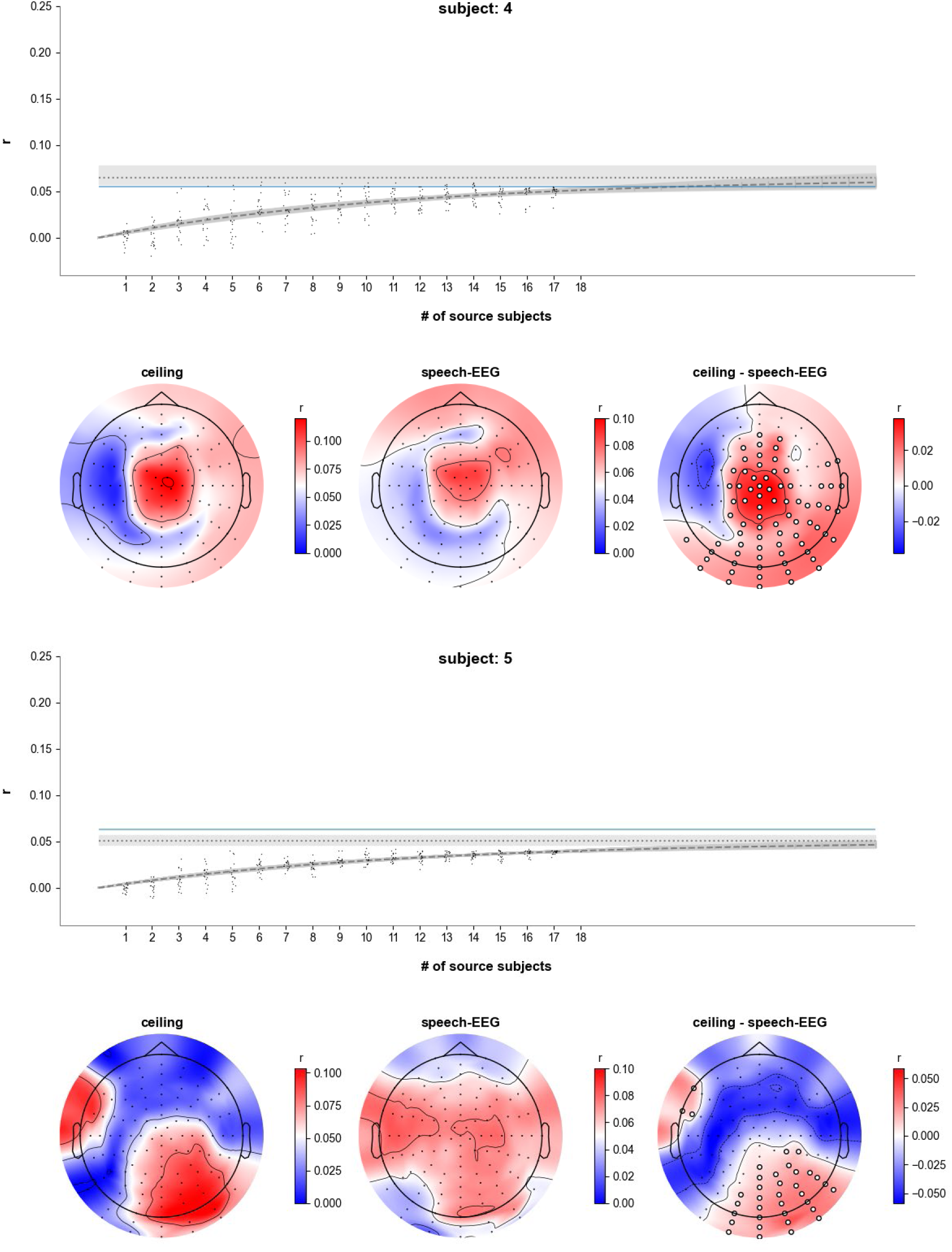
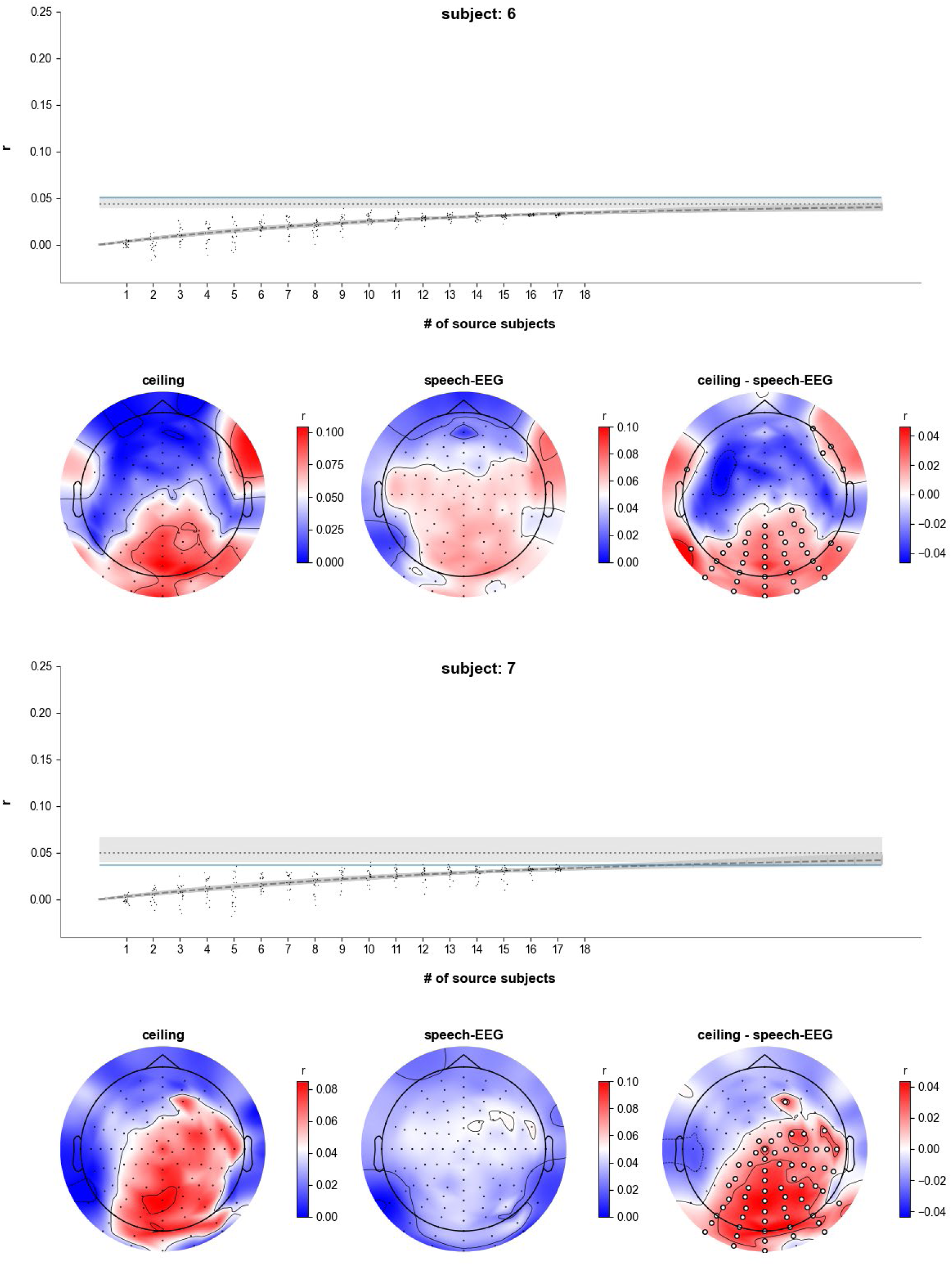
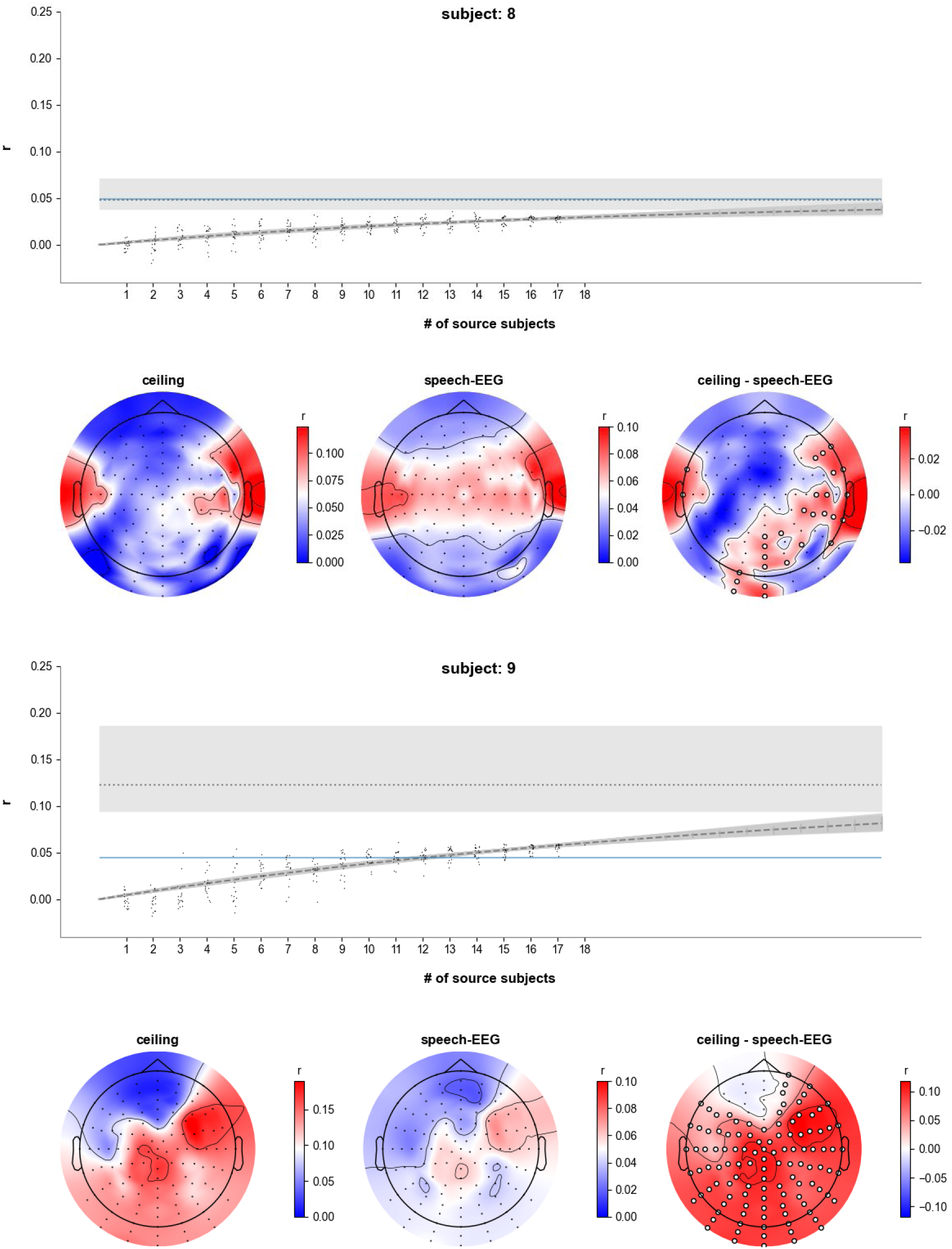
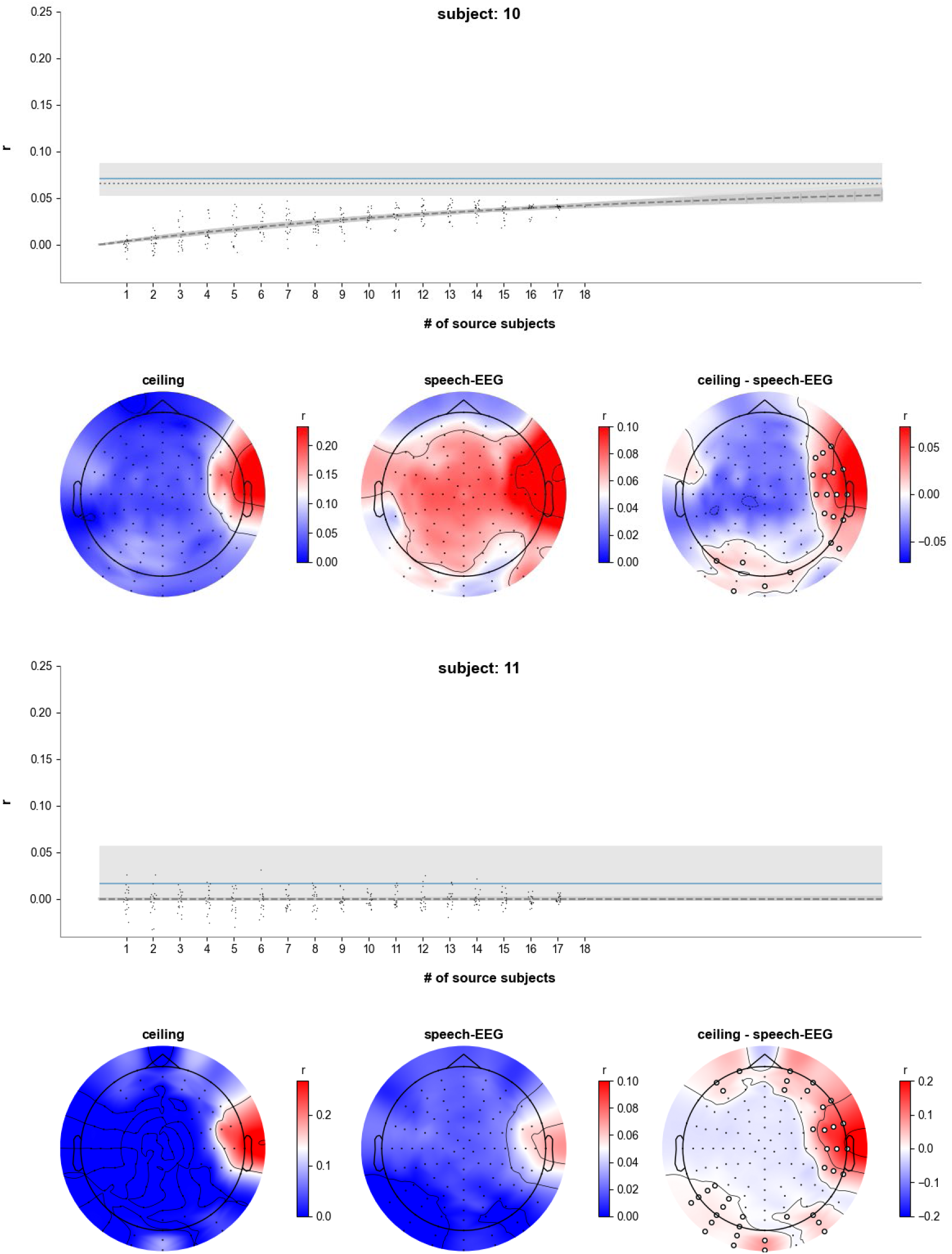
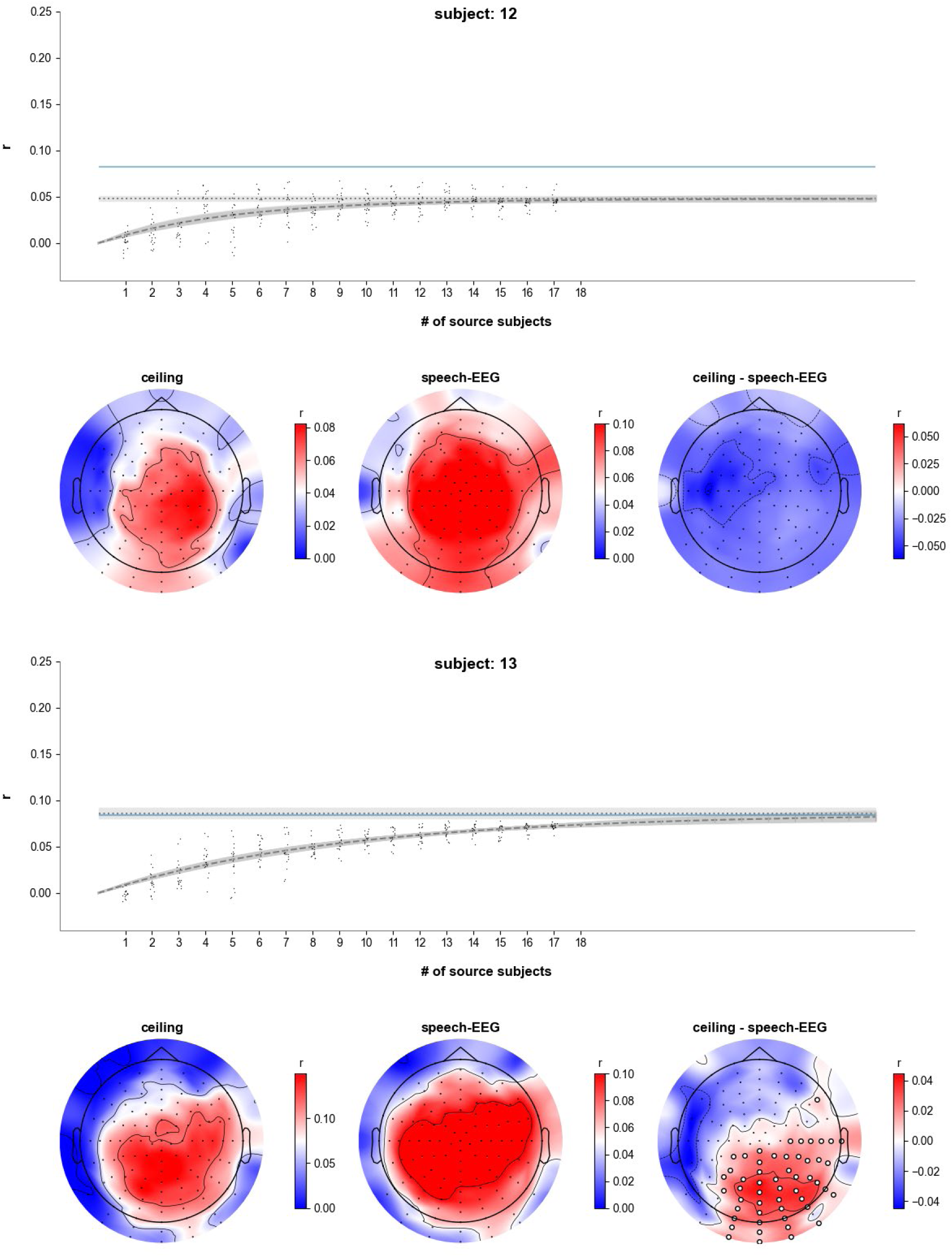
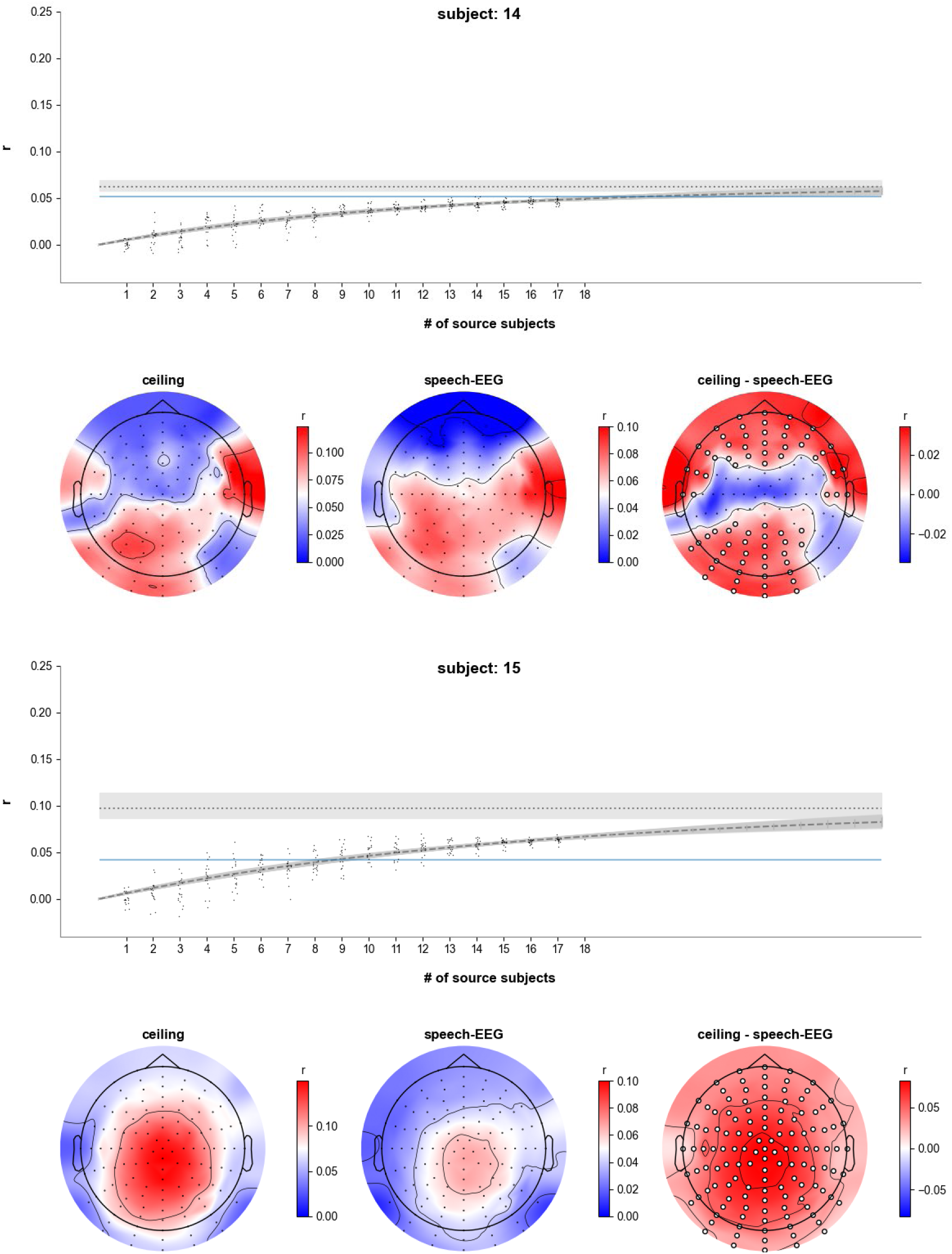
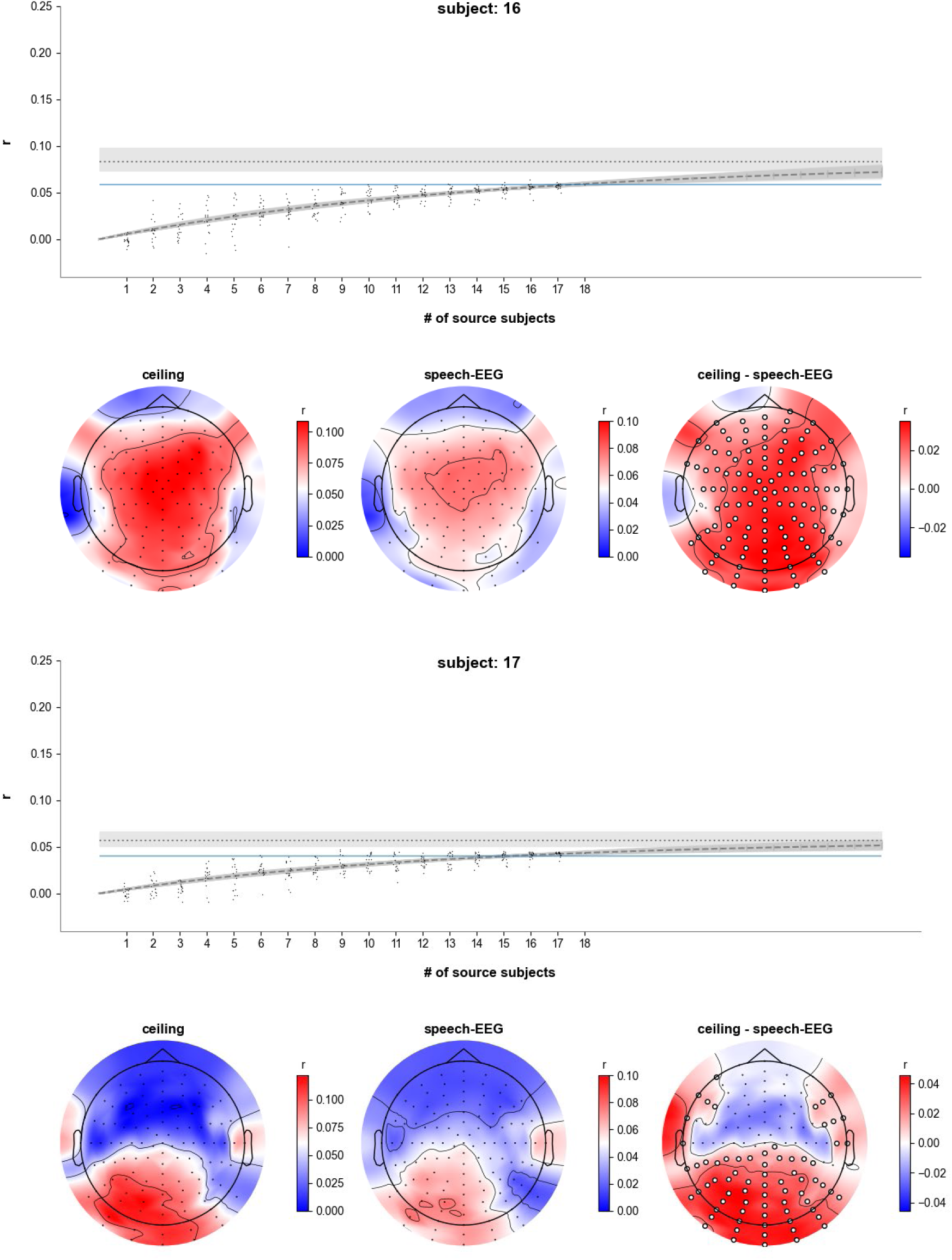
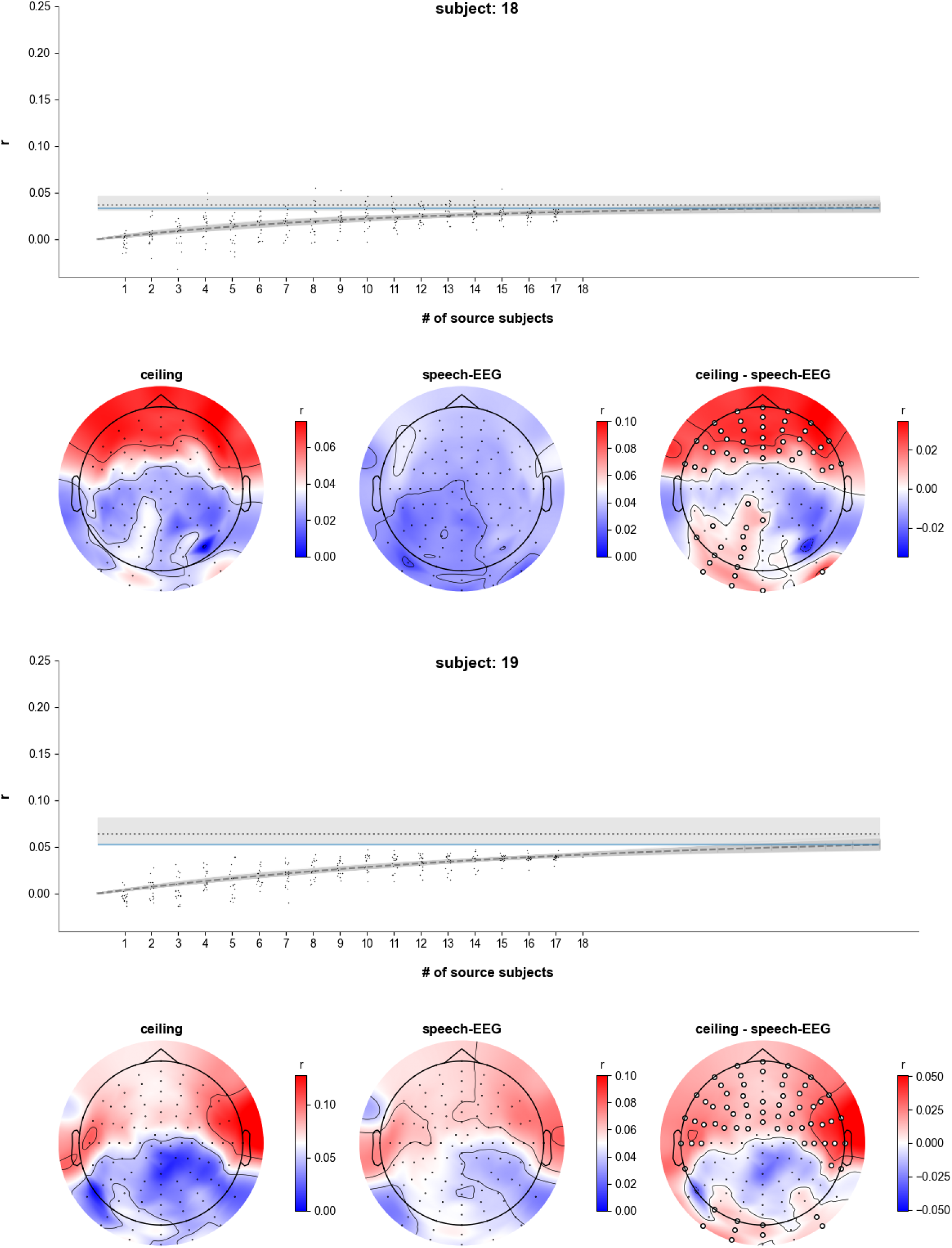
Detailed scalp (upper within each subject’s panel) and channel-level (lower within each subject’s panel) noise ceiling for each subject.

## Footnotes

^1^When discussing the signal-to-noise ratio of EEG it is important to be clear that the “noise” we refer to in this work is not just noise in the traditional sense of line noise or electrical artifacts but also includes all the EEG that is not related to the stimulus. Indeed, most of the “noise” we are talking about in this study – as in many EEG studies – is just EEG signal that is unrelated to the stimulus.

## Notes

### Competing Interest Statement

The authors have declared no competing interest.

